# Structural genomic variation leads to unexpected genetic differentiation in Lake Tanganyika’s sardines

**DOI:** 10.1101/800904

**Authors:** Julian Junker, Jessica A. Rick, Peter B. McIntyre, Ismael Kimirei, Emmanuel A. Sweke, Julieth B. Mosille, Bernhard Wehrli, Christian Dinkel, Salome Mwaiko, Ole Seehausen, Catherine E. Wagner

## Abstract

Identifying patterns in genetic structure and the genetic basis of ecological adaptation is a core goal of evolutionary biology and can inform the management and conservation of species that are vulnerable to population declines exacerbated by climate change. We used reduced representation genomic sequencing methods to gain a better understanding of genetic structure among and within populations of Lake Tanganyika’s two sardine species, *Limnothrissa miodon* and *Stolothrissa tanganicae*. Samples of these ecologically and economically important species were collected across the length of Lake Tanganyika, as well as from nearby Lake Kivu, where *L. miodon* was introduced in 1959. Our results reveal unexpected differentiation within both *S. tanganicae* and *L. miodon* that is not explained by geography. Instead, this genetic differentiation is due to the presence of large sex-specific regions in the genomes of both species, but involving different polymorphic sites in each species. Our results therefore indicate rapidly evolving XY sex determination in the two species. Additionally, we found evidence of a large segregating inversion in *L. miodon*. We found all inversion karyotypes throughout Lake Tanganyika, but the frequencies vary along a north-south gradient, and differ substantially in the introduced Lake Kivu population. We do not find evidence for significant isolation-by-distance, even over the hundreds of kilometers covered by our sampling, but we do find shallow population structure.

## Introduction

Pelagic mixed fish stocks are notoriously difficult to manage (Belgrano & Fowler 2011; Botsford et al. 1997) and part of this challenge lies in identifying Management Units (MUs) which are demographically independent and genetically distinct populations. In a habitat without physical barriers, low genetic differentiation is typical, as there exist few environmental restrictions to gene flow. However, there are increasingly cases detected where small genomic differences lead to important variation in life history, influencing population resilience to fishing pressure (Berg et al. 2017; Hutchinson 2008; Kirubakaran et al. 2016). The use of next generation sequencing methods, which can resolve such fine-scale genetic structure through sampling a large proportion of the genome, is therefore needed to shed light on population structure, particularly in species with low genetic differentiation.

Of particular recent interest is the role of genomic regions with reduced recombination rates, such as chromosomal inversions (e.g. Berg et al. 2017; Christmas et al. 2018; Kirubakaran et al. 2016; Lindtke et al. 2017), sex chromosome regions (Presgraves 2008; Qvarnstrom & Bailey 2009) or both (Connallon et al. 2018; Hooper et al. 2019; Natri et al. 2019) in generating genetic structure within spatially panmictic populations. The reduced recombination rates in such chromosomal regions may enable local adaptation even when gene flow is high (Kirkpatrick & Barton 2006). Furthermore, it appears that these mechanisms for restricted recombination are more prevalent in sympatric than in allopatric species (McGaugh et al 2012; Castiglio et al 2014), and fixation of inversions is faster in lineages with high rates of dispersal and gene flow (Berg *et al*. 2017; Hooper & Price 2015; Martinez *et al*. 2015). These patterns are consistent with theory where chromosomal rearrangements, capturing multiple co-adapted loci, are favored to spread in the presence of gene flow (Berg et al. 2017; Kirkpatrick & Barton 2006). Identifying the genetic basis of ecological adaptation is thus a high priority in evolutionary ecology and can have important implications for population management.

Pelagic habitats allow for high dispersal rates due to the lack of predominant physical barriers. Well known examples of species from pelagic habitats that carry chromosomal inversions or sex-linked genomic differentiation include Atlantic cod (*Gadus moruha*) (Berg *et al*. 2017; Kirubakaran *et al*. 2019; Kirubakaran *et al*. 2016) and Atlantic herring (*Clupea harengus*) (Lamichhaney *et al*. 2017; Martinez Barrio *et al*. 2016; Pettersson *et al*. 2019). In Atlantic cod and herring populations, low genome-wide divergence is interspersed with highly divergent inverted regions. These inversions in cod distinguish between resident and migrating ecotypes (Berg et al. 2017; Kirubakaran et al. 2016) and males and females (Kirubakaran *et al*. 2019), and in herring they separate spring and fall spawners (Lamichhaney *et al*. 2017; Martinez Barrio *et al*. 2016; Pettersson *et al*. 2019). Additionally, inverted genomic regions in sticklebacks are involved in the divergence between lake and stream ecotypes (Marques et al. 2016; Roesti et al. 2015).

Lake Tanganyika is volumetrically the second largest lake in the world, consisting of deep basins in the north (∼1200 m) and south (∼1400 m), and a shallower basin (∼800 m) in the central region (Fig. 1A) (McGlue et al. 2007). At 9-12 million years in age (Cohen et al. 1993), it hosts a long history of evolution, which has produced remarkable animal communities consisting largely of endemic species (Coulter 1991). Among these endemics are two sardine species, *Stolothrissa tanganicae* and *Limnothrissa miodon*, which are sister taxa belonging to monospecific genera. Wilson et al. (2008) showed evidence that the sardines of Lake Tanganyika descend from relatives in western Africa and that these sister taxa diverged from a common ancestor about 8 MYA, presumably within Lake Tanganyika due to the endemic distributions of these species and the age of the basin.

**Figure 1.**
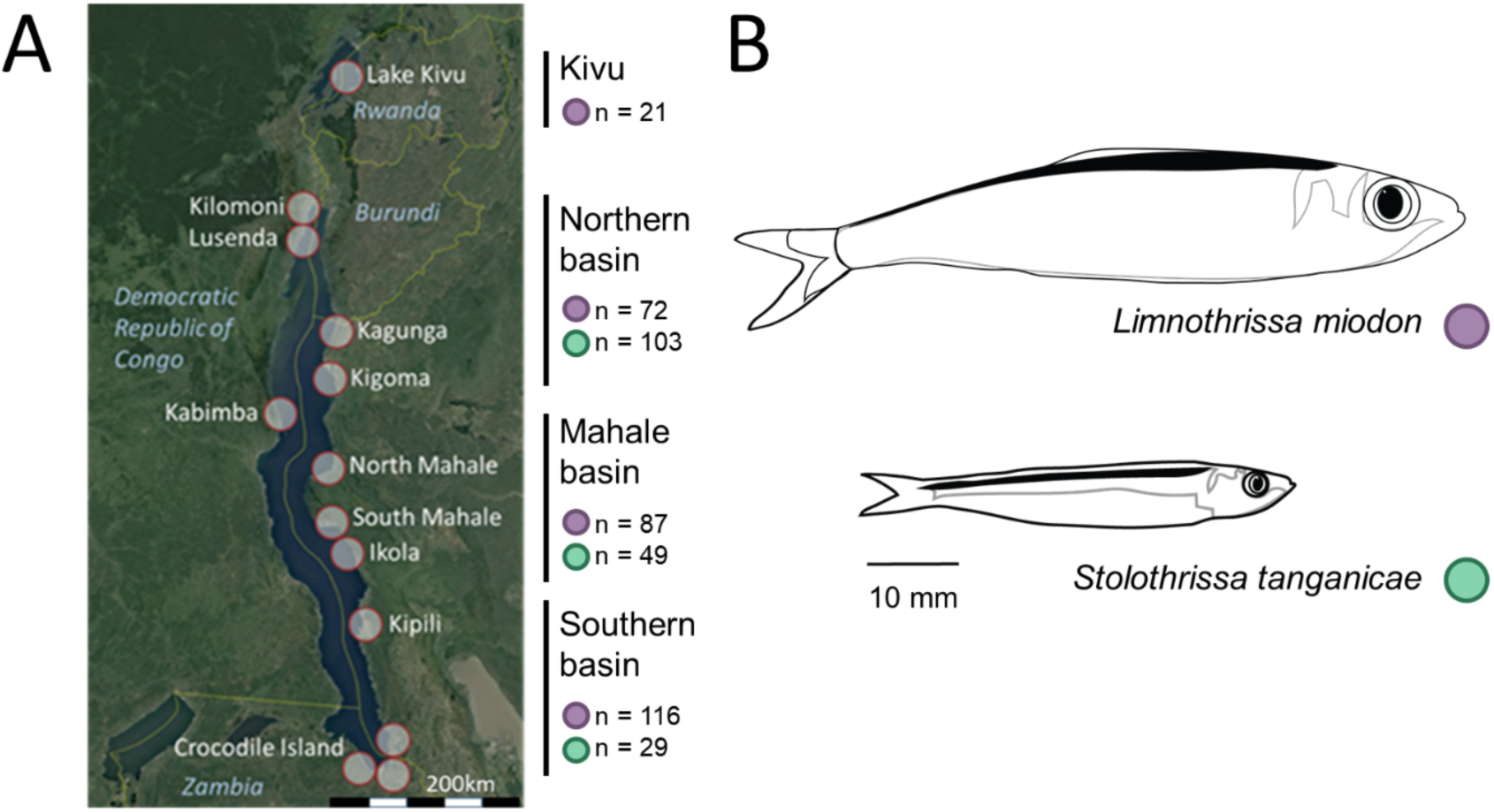
(A) Map of Lake Tanganyika, with sampling sites labeled and number of fish sequenced from the three basins within Lake Tanganyika and Lake Kivu indicated for each species. (B) Drawings of *Limnothrissa miodon* and *Stolothrissa tanganicae*, with scale indicated, show average mature sizes of the two species. Drawings courtesy of Jimena Golcher-Benavides.

The harvest of *S. tanganicae* and *L. miodon* account for up to 65% of all catches within the lake (Coulter 1976, 1991; Mölsä et al. 2002), contributing to the second largest inland fishery on the continent of Africa (FAO 1995). The fishing industry provides employment to an estimated 160,000 (Van der Knaap et al. 2014) to 1 million people (Kimirei et al. 2008) and is an important source of protein to additional millions living on the shores of Lake Tanganyika and further inland (Kimirei et al. 2008; Mölsä et al. 2002; Sarvala et al. 2002; Van der Knaap et al. 2014). Due to human population growth and an increased demand for protein, fishing pressure has increased during the last decades, resulting in a decline of pelagic fish stocks (Coulter 1991; van der Knaap 2013; Van der Knaap et al. 2014; van Zwieten et al. 2002). In addition, long-term decreases in fish population abundance are likely linked to the observed warming of Lake Tanganyika since the early 1900s, and further warming-induced decline in the lake’s productivity is expected during the 21^st^ century (Cohen et al. 2016; O’Reilly et al. 2003; Verburg & Hecky 2003; Verburg et al. 2003). Consequently, there is increasing recognition of the need to develop sustainable management strategies for the lake’s pelagic fish stocks (Kimirei et al. 2008; Mölsä et al. 1999; Mölsä et al. 2002; van der Knaap 2013; Van der Knaap et al. 2014; van Zwieten et al. 2002).

Despite the economic importance of the pelagic fisheries in this lake, little previous work has investigated the diversity and population structure of the key pelagic fish species or their evolutionary origins (but see De Keyzer et al. 2019; Hauser et al. 1995, 1998; Wilson et al. 2008). Lake Tanganyika’s enormous size (∼670km from north to south) harbours the potential for isolation by distance patterns to emerge, and for spatial segregation that may lead to temporal variation in spawning and life history timing between distant sites. The nutrient availability in the water column is regulated by trade winds and complex differential cooling, leading to regions of higher and lower productivity within the lake (Bergamino *et al*. 2010; Plisnier *et al*. 1999; Verburg *et al*. 2011). Mulimbwa et al (2014a and b) found that reproduction may additionally correlate with food availability, suggesting that reproduction may vary spatially in conjunction with spatial differences in primary productivity. Furthermore, spawning peaks in *S. tanganicae* occur at different times in the southern and northern basin of Lake Tanganyika (Chapman & van Well 1978; Ellis 1971). Such patterns of spatial and temporal differences in spawning suggest that geographic differentiation in life history timing may exist between distant sites.

Despite their close relationship, *S. tanganica*e and *L. miodon* have substantial differentiation in life histories. *S. tanganicae* forms large schools and has a fully pelagic life cycle, including pelagic spawning (Coulter 1970, Mannini 1998a). Fertilized embryos of *S. tanganicae* develop while they sink in the water column (at a rate of 4-5 cm/minute) and the larvae hatch after 24-36h (Matthes 1967). There is evidence that juveniles between 10mm and 50mm tend to move in-shore to escape predation, forming mixed schools with *L. miodon* juveniles, and move off-shore again at sizes larger than 50mm (Coulter 1991). In contrast, in *L. miodon* spawning occurs in the near shore (Coulter 1991, Ellis 1971, Mannini 1998a) and individuals only move to the pelagic once they reach large sizes. For *L. miodon* in Lake Kivu, spawning fish have been found both inshore and offshore, so it is unclear whether spawning is strictly littoral in this introduced population (Spliethoff *et al*. 1983).

*S. tanganicae* has a maximum mean total length of about 100mm, compared to *L. miodon* where the adult mean total length is about 120mm (Coulter 1991; Mannini *et al*. 1996), with the former species living about 1.5 years, whereas the latter lives for about 2.5 years (Coulter 1991; Pearce 1985). Sexually mature individuals (*S. tanganica*e: female ∼75mm, males ∼70mm; *L. miodon*: females ∼75 mm and males at ∼64mm in southern Lake Tanganyika and ∼62mm for females and ∼61mm for males in Lake Kivu; Ellis 1971, Spliethoff et al 1983) exist year round but fisheries data indicate that spawning peaks exist (Coulter 1970, Ellis 1971, Mannini 1998a, Marlier 1957), with peaks happening earlier in the southern than in the northern part of the lake in *S. tanganica*e (Coulter 1991, Ellis 1971). In introduced *L. miodon* in Lake Kivu, spawning takes place year round but peaks can also be observed (Spliethoff *et al*. 1983)

Juvenile *S. tanganica*e feed mostly on phytoplankton (Coulter 1991) and switch to zooplankton, shrimp and fish larvae when they move offshore (>50mm length) (Chèné 1975). Juvenile *L. miodon* feed mainly on phyto- and later on zooplankton and shrimp, but larger specimens also prey on *S. tanganicae* or young *L. miodon* (Coulter 1991; Mannini 1998a). In the pelagic, schools are in deep waters during the day and move upwards at dusk and downwards at dawn following the diurnal vertical migration movements of copepods. Although this vertical migration is clear, different opinions exist about the lateral migration of the two species. There are indications for extensive movement based on echo sounding studies by Johannesson (1975) and Chapman (1976), and van Zwieten et al. (2002) suggest in- and offshore movement based on catch data (Coulter 1991). In contrast, other studies have suggested little movement in *S. tanganica*e due to local increases in *S. tanganica*e populations when local predator abundance declined due to fishing pressure at the same locality (Coulter 1991, Ellis 1978).

There are some indications of genetic differentiation within pelagic fish populations of Lake Tanganyika known from basic genetic work conducted two decades ago. For the sardines, these studies found no clear genetic population structure at a large geographical scale (Hauser et al. 1998; Kuusipalo 1999), but some small-scale differences were found for *L. miodon* (Hauser et al. 1998). However, the genetic methods used in these older studies (RAPDs and microsatellites) have limited power and are known to suffer from error (RAPD, Williams et al. 1990). In *S. tanganicae*, De Keyzer et al. (2019) recently used an mtDNA data set and a restriction site associated DNA (RAD) sequencing data set based on 3504 SNPs and 83 individuals, sampled from the north, middle, and south of Lake Tanganyika, finding little evidence for spatial genetic structure.

In this study, we focus on analysing patterns of genetic diversity and divergence in both sardine species, *S. tanganicae* and *L. miodon*, using next-generation sequencing based approaches. We sampled sardines from 13 sites spanning from the north to the south of Lake Tanganyika (Fig. 1). We also included *L. miodon* individuals from the introduced population of this species present in Lake Kivu. Our null hypothesis was simple: the surface water of a large lake is horizontally well mixed and therefore provides a relatively homogeneous habitat. Pelagic fish can move freely and therefore due to the uniform environment, we should expect a lack of genetic structure of their populations due to free interbreeding. Using reduced representation genomic sequencing (RAD, Baird et al. 2008) we indeed do not find substantial spatial genetic structure in either species, supporting this null hypothesis. However, many loci deviating from Hardy-Weinberg equilibrium differentiated the sexes in our samples, suggesting that these species have large sex-determining regions. Furthermore, we find additional cryptic genetic diversity in *L. miodon* that is consistent with the existence of a chromosomal inversion. Additionally, there is some evidence for very weak yet distinct sympatric genetic groups in both sardines which differ in frequency across sampling sites but are overlapping. However, these signals are based on few loci, and this structure is not evident in PCAs. This weak genetic structure may be the result of selection, or additional structural genomic variation. However, there is no evidence for significant isolation-by-distance, despite the large geographic scale at which we sampled. The low spatial genetic structure within these species facilitated the detection of these differentiated loci and genomic structural variation, which may be related to sex-specific and local adaptation.

## Material and Methods

### Study system and sampling

Our samples from Lake Tanganyika come from Tanzanian, Congolese and Zambian sites and were collected between the years 2015 and 2017. Additionally, we added Rwandan *L. miodon* sampled in 2013 from Lake Kivu, where the species was introduced during the 1950s (Collart 1960, 1989; Hauser *et al*. 1995) (Fig. 1 and Table 1). These fish included some individuals that were collected live, some that were collected dead (from fishermen), and some that were collected dried (from markets). For fish collected live, each fish was processed according to our standard sampling protocols, during which we took a cuvette photograph of the live fish and subsequently euthanized the fish with an overdose of MS222, and took fin clips and muscle tissue samples for genetic analysis (stored in ethanol) and stable isotope analysis (dried), respectively. The specimens were preserved in formaldehyde and then archived in the collections at EAWAG (samples from the years 2013, 2016, 2017), the University of Wyoming Museum of Vertebrates (2015 samples), or the University of Wisconsin-Madison (2015 samples). Most fish for this study were obtained from fishermen and were already dead, and in this case we completed this same protocol without euthanasia. For fish that were desiccated prior to collection, we stored the whole fish dried, took desiccated fin clips for DNA extraction, and preserved the remaining dried specimen in museum collections.

**Table 1.** Fish collected and sequenced from Democratic Republic of Congo (DRC), Tanzania (TNZ), Zambia (ZM) and Rwanda (RW).

| **Number of sequenced individuals** |  |  |  |  |  |
| --- | --- | --- | --- | --- | --- |
| *Stolothrissa*<br>*tanganyicae* | *Limnothrissa*<br>*miodon* | Site | Basin | Country | Sampling year |
| 0 | 21 | Lake Kivu | ----- | RW | 2013 |
| 7 | 0 | Kilomoni | North | DRC | 2016 |
| 15 | 0 | Lusenda | North | DRC | 2016 |
| 15 | 20 | Kagunga | North | TNZ | 2017 |
| 61 | 50 | Kigoma | North | TNZ | 2015, 2016, 2017 |
| 5 | 2 | Kabimba | North | DRC | 2016 |
| 25 | 37 | North Mahale | Mid | TNZ | 2015, 2016 |
| 6 | 38 | South Mahale | Mid | TNZ | 2015, 2016 |
| 18 | 12 | Ikola | Mid | TNZ | 2017 |
| 12 | 61 | Kipili | South | TNZ | 2017 |
| 0 | 41 | Kasanga | South | TNZ | 2017 |
| 17 | 1 | Mbete | South | ZM | 2016 |
| 0 | 13 | Crocodile Island | South | ZM | 2016 |
| **181 Samples** | **296 Samples** | **13 Sites** | **3 Basins** | **4 Countries** | **4 Years of Sampling** |

### Phenotypic sexing

Tanganyikan sardines caught by fishermen are frequently dried after landing and although this does not inhibit the extraction of high-quality DNA, desiccated individuals cannot be accurately sexed. Therefore, we dissected 34 *L. miodon* and 15 *S. tanganicae* that were euthanized and preserved in formalin just after being caught. We chose only individuals that were > 70mm in length, fully mature, and in excellent condition, to accurately determine whether each fish, based on their gonads, was male or female. We used these phenotypically sexed individuals to test whether inferred genetic groups correlated to sex in each species.

### RAD sequencing

Once returned from the field, tissues for genetic analysis were stored in ethanol at -20°C prior to DNA extraction. We extracted DNA from 475 individuals (181 *S. tanganicae;* 294 *L. miodon*) and obtained genomic sequence data for these individuals using a reduced-representation genomic sequencing approach (RADseq, (Baird *et al*. 2008)). The DNA from all individuals was extracted using Qiagen DNeasy Blood and Tissue kits (Qiagen, Switzerland). All individuals were barcoded, then pooled and divided into 10 RAD libraries for sequencing. For 190 individuals collected in 2015, the DNA was standardized to 20ng/µL at the University of Wyoming, prepared for RAD sequencing by Floragenex Inc. (Eugene, Oregon), and sequenced at the University of Oregon on an Illumina HiSeq2000 (100bp SE), with one library sequenced per lane. For the Floragenex libraries, library prep followed the protocol in Baird *et al*. (2008), and individuals were multiplexed in groups of 95 individuals using P1 adapters with custom 10 base pair barcodes, and fragments between 200 and 400bp were selected for sequencing. To avoid library effects, each individual was sequenced in two different libraries and the reads were combined after sequencing. The other 296 individuals collected in 2013, 2016 and 2017, were prepared for sequencing at EAWAG following the protocol by Baird *et al*. (2008) with slight modifications, including using between 400ng and 1000ng genomic DNA per sample and digesting with *SbfI* overnight. We multiplexed between 24 and 67 of these individuals per library and used P1 adapters (synthesized by Microsynth) with custom six to eight base pair barcodes. These six libraries were sheared using an S220 series Adaptive Focused Acoustic (AFA) ultra-sonicator (Covaris Inc. 2012) with the manufacturer’s settings for a 500 bp mean fragment size. The enrichment step of library preparation was done in eight aliquots with a total volume of 200 μl. Volumes were combined prior to a final size selection step using a SageELF (Sage Scientific Electrophoretic Lateral Fractionator; Sage Science, Beverly, MA), during which we selected fragments with a size between 300 and 700bp. Sequencing was done by the Lausanne Genomic Technologies sequencing facilities of the University of Lausanne, Switzerland. We sequenced each of six libraries on a single lane of an Illumina HiSeq2000 (100bp SE).

### Sequence data preparation

We filtered raw sequencing reads from each library to remove common contaminants by first removing PhiX reads using bowtie2 (Langmead & Salzberg 2012), and then filtering reads for an intact *SbfI* restriction site. We then de-multiplexed the fastq file and trimmed the reads down to 84 nucleotides using process_radtags from Stacks v1.26 (Catchen *et al*. 2013) and a custom bash script. The FASTX-toolkit v.0.0.13 (http://hannonlab.cshl.edu/fastx_toolkit/) was used for quality filtering. In a first step, we kept only reads with all base quality scores greater than 10; in a second step, we removed all reads with more than 5% of the bases with quality score below 30.

### Reference genome assembly

We generated a reference genome from a male *L. miodon* individual collected near Kigoma, Tanzania, in 2018, to use in aligning our RAD sequencing reads. High molecular weight DNA was extracted from fin tissue using the Qiagen HMW gDNA MagAttract Kit, and then libraries were prepared using 10X Genomics Chromium library preparation at the Hudson-Alpha Institute for Biotechnology Genomic Services Laboratory (Huntsville, AL). The sequencing libraries were then sequenced on the Illumina HiSeq Xten platform (150bp PE reads). Read quality was checked using FASTQC (v 0.1.2, Andrews 2010), and reads were then assembled using 10X Genomics’ Supernova v2.0 assembly software, using a maximum of 500 million reads. We assessed assembly completeness using QUAST-LG (v 5.0.0, Mikheenko *et al*. 2018), which computes both standard summary statistics and detects the presence of orthologous gene sequences.

### Alignment to the reference genome and SNP calling

Reads for all *L. miodon* and *S. tanganicae* individuals were aligned to the reference genome using BWA mem (v0.7.17, Li & Durbin 2009) with default settings, after the initial read filtering steps with the FASTX-toolkit discussed above. We chose to align individuals from both species to our draft reference genome after observing high mapping rates in both species. Following alignment, we excluded any individuals with < 50,000 reads aligned to the reference genome. Subsequent analyses were performed on the remaining 178 *S. tanganicae* and 287 *L. miodon* individuals with greater than 50,000 reads aligned.

We identified variable sites in three different sets of individuals using SAMtools mpileup (v1.8, Li et al. 2009b) and bcftools (v1.8, Li *et al*. 2009a): (1) all individuals; (2) only *L. miodon* individuals; and (3) only *S. tanganicae* individuals. In each of these data sets, we omitted indels and kept only high-quality biallelic variant sites (QUAL < 20 and GQ > 9). We obtained consistent results using different combinations of more stringent and relaxed filtering steps. The results shown here are based on a filtering as follows: within the species-specific data sets (where we had either only *S. tanganicae* or only *L. miodon* individuals), we filtered SNPs using VCFTOOLS (Danecek *et al*. 2011) to allow no more than 50% missing data per site, removed SNPs with a minor allele frequency less than 0.01, and only called genotypes with a minimum read depth of 2. For the data set including both species, we relaxed the missing data filter to allow sites with up to 75% missing data.

We checked for library effects within our data by plotting a PCA of genotypes called within *L. miodon* and within *S. tanganicae*, to ensure that our data from different years and library preparation methods were compatible. After observing evidence for library effects within both our *L. miodon* and *S. tanganicae* data sets (Fig. S1A and S1C), we filtered to keep only SNPs present in individuals from libraries sequenced at both facilities. After removing these library-specific SNPs, we again checked for library effects to ensure that they no longer were evident (Fig. S1B, S1D, S2 and S3).

### Population structure and outlier detection

We used the species-combined data set for two analysis: first, for a PCA testing for genetic differentiation between *S. tanganicae* and *L. miodon*; and second, to test if sex linked loci overlap between the two species. All other analyses were done using the single species data sets.

After removing individuals with more than 25% missing data at the genotyped SNPs, we used the species-combined data set to conduct principal component analysis (PCA) on the genotype covariance matrix, using prcomp from the package stats (v3.5.3) in R. To delineate and visualize distinct groups within our data without a priori group assignments, we performed genetic-based K-means clustering (find.clusters from adegenet; Jombart 2008) for K=4 groups. We then used these groupings to assign individual fish to species and clusters within species. We combined these clustering results with sexed phenotypes to confirm the identity of each of the four clusters. We then conducted two discriminant analysis of principal components (DAPC, Jombart *et al*. 2010) analyses to identify loci contributing to the difference between the two clusters within each of the species. To identify loci with significant loadings, we simulated null expectations by randomizing genotypes among individuals at each locus, performing DAPC on the randomized data sets, and repeating the randomization 1000 times. We used the loadings from these randomized DAPC runs to create null distributions for each of *L. miodon* and *S. tanganicae*. We identified SNPs with loadings above the 99% quantile of the null distribution as significant in each analysis, and compared those SNPs identified in each species.

We then moved to working with each species individually in the species-specific data sets. We first used PCA on the genotype covariance matrices to visualize structure within each species. After observing that the primary axis of differentiation in both *S. tanganicae* and *L. miodon* was based on sex in the species-combined data set, we then used the single-species SNP data sets and the R package adegenet (Jombart 2008) to conduct DAPC on males versus females of each species to identify loci contributing to these sex differences. We simulated null expectations for SNP loadings in the DAPC as described above, for each of *L. miodon* and *S. tanganicae*. We again identified SNPs with loadings above the 99% quantile of the null distribution as significant in each analysis. We then calculated heterozygosity for these sex-associated loci using adegenet in R.

In *L. miodon*, the secondary axis of genetic differentiation clearly split the population into three distinct genetic groups. To investigate the genetic basis of these groupings, we again used genetic-based K-means clustering to assign individuals to groups. After assigning all individuals to one of the three distinct groups based on clustering, we calculated the frequencies of the three groups at each sampling site and within each lake basin (i.e. north, middle, south). To determine whether the distribution of individuals among the clusters varied between regions in Lake Tanganyika, we conducted a two-proportion z-test (prop.test in R) between the three general regions in Lake Tanganyika, as well as between each of these and Lake Kivu.

We then used DAPC to identify the loci with high loadings on the differentiation between the two most extreme groups, using the *L. miodon*-only data set with Lake Kivu individuals omitted. Once again, we identified SNPs significantly associated with group delineation by creating a null distribution and selecting those SNPs above the 99% quantile of the null. We then calculated heterozygosity for these significant loci using adegenet in R. Because patterns of heterozygosity were consistent with these three groups being determined by a segregating chromosomal inversion, we then tested whether the three genotypes are in Hardy-Weinberg Equilibrium within each of the three distinct geographic regions and within Lake Kivu.

*S. tanganicae* and *L. miodon* are sister species and if they share sex determining regions, then we would expect them to map to similar locations on the *L. miodon* reference genome. However, if the sets of sex-linked SNPs of each species map to different regions, then we expect that one or either of the species have switched the chromosomes used in sex determination (“turnover” of sex determining regions). This could occur either from the translocation of a sex-determining locus to a new genomic location, or due to the origin of a new mutation with sex-determination function (Jeffries *et al*. 2018). We therefore assessed whether the same genomic regions explain genetic differentiation between sexes in the two species. For this, we compared the location of SNPs identified in each of the *S. tanganicae* and *L. miodon* DAPC analyses, both using the species-specific and combined SNP data sets, by calculating the proportion of scaffolds shared among the two sets of significant SNPs. As an additional comparison between the two species, we calculated the proportion of *L. miodon* sex-linked SNPs that were polymorphic in *S. tanganicae*, and vice versa, as well as the observed heterozygosity of *L. miodon* individuals at *S. tanganicae* sex-linked SNPs, and vice versa.

### Geographic population structure

For each species separately, we investigated population structure beyond sex differences to determine whether there is any geographic signal of differentiation within the two species. We removed the scaffolds containing sex-associated SNPs in the species-specific data sets for each species. In *L. miodon* we additionally removed the scaffolds containing SNPs associated with the inverted region. We then calculated F_ST_ between all sampling site pairs and between basins using the Reich-Patterson F_ST_ estimator (Reich *et al*. 2009) and estimated 95% confidence intervals using 100 bootstrap replicates. We used these estimates in a Mantel test (mantel.randtest from adegenet in R) for each species, to test for a possible association between genetic distances and Euclidean geographic distances between sites (i.e. isolation by distance). For the Mantel tests, we calculated F_ST_/(1-F_ST_) and omitted Lake Kivu, as well as locations with fewer than 10 individuals.

To formally test for structure within the *L. miodon* and *S. tanganicae* populations, we used the hierarchical Bayesian genetic-based clustering program entropy (Gompert et al. 2014), a program and model much like STRUCTURE (Pritchard et al. 2000, Falush et al. 2003), which leads to estimates of allele frequencies in putative ancestral clusters and admixture proportions for individuals. Both entropy and STRUCTURE incorporate no *a priori* assumptions about assignment of individuals to clusters and require only the specification of the number of ancestral clusters (K). Entropy additionally incorporates uncertainty about individuals’ true genotypes by taking genotype likelihoods from bcftools (Li 2011) as input. Thus, the model integrates over genotype uncertainty, appropriately propagating uncertainty to higher levels of the model. Additionally, entropy uses calculations of deviance information criterion (DIC) for model fit to choose among models with different numbers of ancestral population clusters (K).

To compare support for different numbers of clusters, we ran entropy for K=1 to K=10 for each species. After removing SNPs on scaffolds containing sex-associated SNPs (and removing inversion-related scaffolds in *L. miodon*), we ran three independent 80,000 MCMC step chains of entropy for each value of K, discarding the first 10,000 steps as burn-in. We retained every 10^th^ value (thin=10) and obtained 7000 samples from the posterior distribution of each chain. We estimated posterior means, medians, and 95% credible intervals for parameters of interest. We checked MCMC chains for mixing and convergence of parameter estimates by plotting a trace of the MCMC steps for parameters of interest. We then calculated DIC for each value of K and used these to assess which model provided the best fit for the structure in our data. Finally, we conducted an ANOVA on group assignment probabilities (q) for individuals, using sampling site as a factor, to determine whether the means of assignment probabilities to each group differed significantly between sites. We additionally assigned individuals to groups for K > 1 using a threshold of group membership probability of q = 0.6, and calculated Reich-Patterson F_ST_.

### Genetic diversity within clusters

We calculated genetic diversity within the different intraspecific groups using the aligned BAM files in ANGSD (v0.931, Korneliussen *et al*. 2014), again using the *L. miodon* genome as a reference. Methods employed in ANGSD take genotype uncertainty into account instead of basing analyses on called genotypes, which is especially useful for low- and medium-depth genomic data (Korneliussen *et al*. 2014), such as those obtained using RAD methods. From these alignment files, we first calculated the site allele frequency likelihoods based on individual genotype likelihoods (option -doSaf 1) using the samtools model (option -GL 1), with major and minor alleles inferred from genotype likelihoods (option -doMajorMinor 1) and allele frequencies estimated according to the major allele (option - doMaf 2). We filtered sites for a minimum read depth of 1 and a maximum depth of 100, minimum mapping quality of 20, and minimum quality (q-score) of 20. In addition, we omitted all scaffolds that contained sex or inversion loci. From the site allele frequency spectrum, we then calculated the maximum likelihood estimate of the folded site frequency spectrum (SFS) using the ANGSD realSFS program (with option -fold 1). The folded SFS was used to calculate per-site theta statistics and genome-wide summary statistics, including genetic diversity, using the ANGSD thetaStat program (Korneliussen *et al*. 2013). We performed each of these steps on all fish from each of *L. miodon* and *S. tanganicae*, and then individually for each sampling site within each species.

### Linkage disequilibrium among loci

To investigate the extent to which the loci identified by DAPC are linked to one another, we used PLINK (v1.9, Purcell *et al*. 2007) to calculate pairwise linkage disequilibrium between all pairs of SNP loci in our *L. miodon* and *S. tanganicae* data sets. Linkage disequilibrium was measured as the squared allelic correlation (R^2^, Pritchard & Przeworski 2001). We then subsetted each of these comparisons to include only the sex-linked SNPs identified using DAPC, and compared the distribution of linkage values among the sex-linked SNPs to those values between all SNPs in the data set for each of the two species. We then performed the same comparison for loci implicated in differences among the three inversion groups in *L. miodon*. To determine whether sex and grouping loci are more linked than average across the genome, we performed a Mann-Whitney U test (wilcox.test in R) on the sets of linkage values.

## Results

### Genome assembly and variant calling

The final assembly of the 10X Genomics Chromium-generated reference genome for *L. miodon*, based on ∼56x coverage, comprised 6730 scaffolds of length greater than 10Kb. The assembly had a scaffold N50 of 456Kb and a total assembly size of 551.1Mb. The BUSCO score of the genome was 83.5% complete single-copy BUSCO genes, 4.62% fragmented and 11.82% missing BUSCO genes. We retained only scaffolds > 10Kb in length for the reference genome used for downstream alignment of the RAD reads.

The Floragenex RAD libraries yielded between 306 and 328 million reads including 21–23% bacteriophage PhiX genomic DNA, while the libraries sequenced at the Lausanne Genomic Technologies sequencing facilities yielded between 167 and 248 million reads each. This resulted in an average of 2.5 million reads per *S. tanganicae* individual and 4.3 million reads per individual in *L. miodon*. On average, the mapping rate for *S. tanganicae* individuals’ RAD reads to the *L. miodon* reference genome was 80.2%, whereas it was 80.0% for *L. miodon* individuals. Mean read depth was slightly lower overall in *S. tanganicae* than *L. miodon* (64.5 reads vs 73.4 reads), and *S. tanganicae* males had slightly lower mean read depths (average 61.0 reads) than *S. tanganicae* females (mean 67.9 reads; Fig. S4). The ratio of read depths between males and females averaged 1.0 in *L. miodon*, while females in *S. tanganicae* averaged 1.12x more reads than males (Fig. S5A). These mapping rates resulted in 1,224,115 unfiltered variable sites in *L. miodon* and 636,238 unfiltered variants in *S. tanganicae*. We removed 6 *S. tanganicae* individuals and 8 *L. miodon* individuals due to too few reads mapped or too much missing data. After filtering for missing data (50% for species specific data sets and 75% for the species combined data set) and minor allele frequency (MAF > 0.01), our species-specific RAD data sets contained 16,260 SNPs from 175 *S. tanganicae* samples and 28,500 SNPs from 288 *L. miodon* samples. The data set for the combined species approach contained 35,966 SNPs. Due to evidence for library effects (Fig. S1), we further removed 7,195 library-specific SNPs in *S. tanganicae* and 10,072 library-specific SNPs in *L. miodon*, resulting in final data sets of 9,065 SNPs for S. *tanganicae* and 18,428 SNPs for *L. miodon*.

### Evidence for distinct sex loci

Principal component analysis revealed two distinct genetic clusters in each species (Fig. 2A). These clusters correspond to sexes identified through sexing of individuals by dissection (n = 14 *S. tanganicae* and 45 *L. miodon* individuals; Fig. 2A and Table S1), and F_ST_ values indicated relatively large genetic differentiation between sexes in both species (Fig. 2B; male-female F_ST_ = 0.097 for *S. tanganicae*, Fig. 2C; F_ST_ = 0.035 for *<u>L.miodon</u>*). Our phenotypic sexing of well-preserved, sexually mature specimens identified seven *S. tanganicae* individuals as female and seven as male, with one individual identified phenotypically as male clustering with the other females genetically (Fig. 2A, Table S1). This individual was likely an immature female. In *L. miodon*, we identified 27 individuals phenotypically as females and 18 as males, all of which were consistent with genetic groups (Fig. 2A, Table S1).

**Figure 2.**
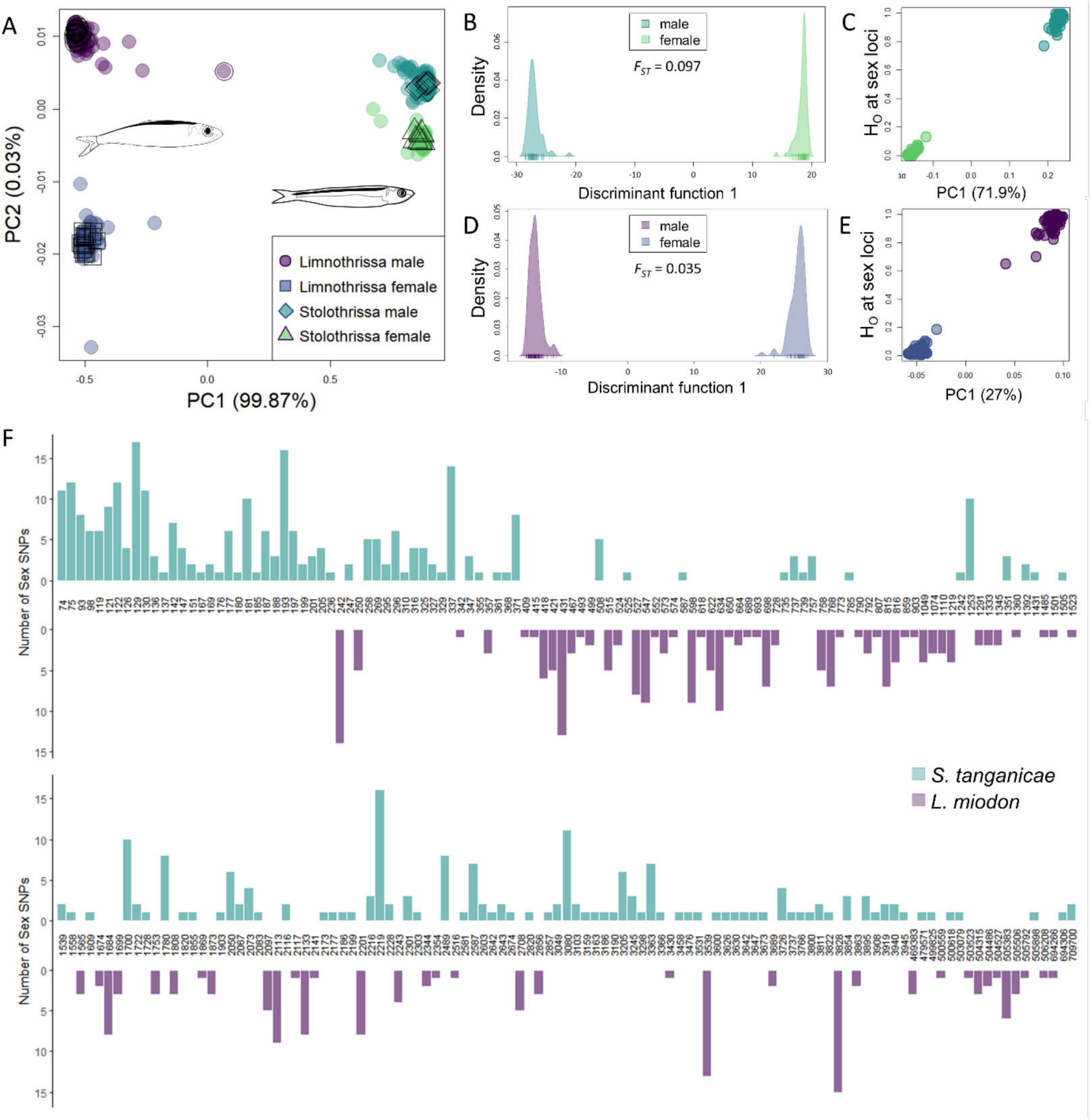
(A) Principal component analysis of all *S. tanganicae* and *L. miodon* individuals combined, colored by species identity and sex. Empty shapes denote individuals that were dissected and for whom sex was determined phenotypically. These dissection phenotypes group into genetic clusters, and therefore were used to identify the sex of each of the genetic clusters. In the combined PCA, the first axis corresponds to species, while the second axis corresponds to sex. Discriminant analysis of principal components (DAPC) results for (B) *S. tanganicae* and (D) *L. miodon* individually demonstrate distinct separation among males and females, with intraspecific differentiation (F_ST_) between the two groups indicated. DAPC on individual species was used to identify loci associated with this differentiation (see Supplementary Figure S1 and S2); observed heterozygosity (H_obs_) of each individual at those loci with high loadings is plotted against the first intraspecific PCA axis in (C) *S. tanganicae* and (E) *L. miodon*, demonstrating both that sex dictates the first axis of differentiation in both species, and that males are the heterogametic sex at these loci in both species. There were 502 significant SNPs differentiating the sexes in *S. tanganicae* and 308 significant SNPs in *L. miodon*, with no overlap between the two species. (F) Distribution of significant sex-associated SNPs in *S. tanganicae* and *L. miodon*, with scaffolds ordered from longest to shortest (and only scaffolds with sex SNPs included). Bars indicate the number of significantly sex-associated SNPs on the given scaffold for the given species, demonstrating that no scaffolds were implicated in sex differentiation in both *S. tanganicae* and *L. miodon*. Scaffold names are indicated across the x-axis. Note: this barplot has been wrapped onto two lines for visual ease and continues from the first line to the second.

In a DAPC to identify the loci underlying the strong genetic differentiation of the sexes for *S. tanganicae*, we selected a loadings cut off of 0.0015 on PC1 based on the null distribution of loadings, which resulted in a total of 502 (5.5%) significant SNPs distributed over 129 scaffolds with high loadings on sex differences (Fig. S6A). In *L. miodon*, we selected a cut-off of 0.00077 on PC1. This cut-off resulted in 308 (1.7%) SNPs across 86 scaffolds with high loadings on sex differences (Fig. S6B). All of these loci show an excess of homozygosity in females and an excess of heterozygosity in males (Fig. 2C and 2E), and no SNPs were significant in both species. In addition, the scaffolds on which these loci were located were non-overlapping between the species (Fig 2F). There were no systematic differences in read depth between males and females at sex-associated SNPs or across scaffolds containing sex-associated SNPs in either species (Fig. S5B). In addition, there was no systematic difference in mean read depth between species or sexes on scaffolds containing sex-associated SNPs (Fig. S5C). The scaffolds containing these sex-associated SNPs span 237Mb of the reference genome for *S. tanganicae* (43.0% of the reference assembly) and 76.6Mb in *L. miodon* (13.9% of the reference assembly).

To test if the sex-linked loci overlap between the species, we used the species-combined data set to perform DAPC between sexes for each species individually and identified loci with high loadings. Using this approach, we identified 570 SNPs across 133 scaffolds in *S. tanganicae* (loading > 0.0006) (Fig. S7A) and 334 SNPs across 91 scaffolds in *L. miodon* linked to sex (loading > 0.001) (Fig. S7B). These two sets of loci were again completely non-overlapping, suggesting that the sex-linked loci are unique in each species. In addition, the scaffolds on which these loci were located were again non-overlapping between the species (Fig. S7A and S7B). When examining *S. tanganicae* sex-linked SNPs in *L. miodon* individuals, only 2.5% are polymorphic, and only 0.8% of *L. miodon* sex-linked SNPs are polymorphic in *S. tanganicae*. In addition, the sex loci for each species do not show the same patterns of heterozygosity in the other species (Fig. S8).

### Evidence for a segregating inversion in L. miodon

*L. miodon* from Lake Kivu are divergent from individuals in Lake Tanganyika, but this differentiation is weaker than that between the three groups observed within Lake Tanganyika (Fig. 3C, 4A). The *L. miodon* individuals from Lake Kivu form additional clusters that are distinct from, but parallel to, the Tanganyika clusters along the second and third PC axis (Fig. 3C, 4A). Within Lake Tanganyika, we found individuals of all three clusters at single sampling sites, and there is no clear geographic signal to these groups (Fig. 3C). DAPC analysis of the two most differentiated groups within Lake Tanganyika identified 91 SNPs across 27 scaffolds with high loadings contributing to group differences (> 0.00077; Fig. 4B, 4C and Fig. S9). Among these SNPs with high loadings, we found that two clusters of *L. miodon* individuals were predominantly homozygous for opposite alleles, while the third group consisted of heterozygotes at these loci (Fig. 4D). This suggests that the three distinct genetic groups we observe result from a segregating inversion, with two of the groups representing homokaryotypes and the third a heterokaryotype for these SNPs (Fig 4D and S9).

**Figure 3.**
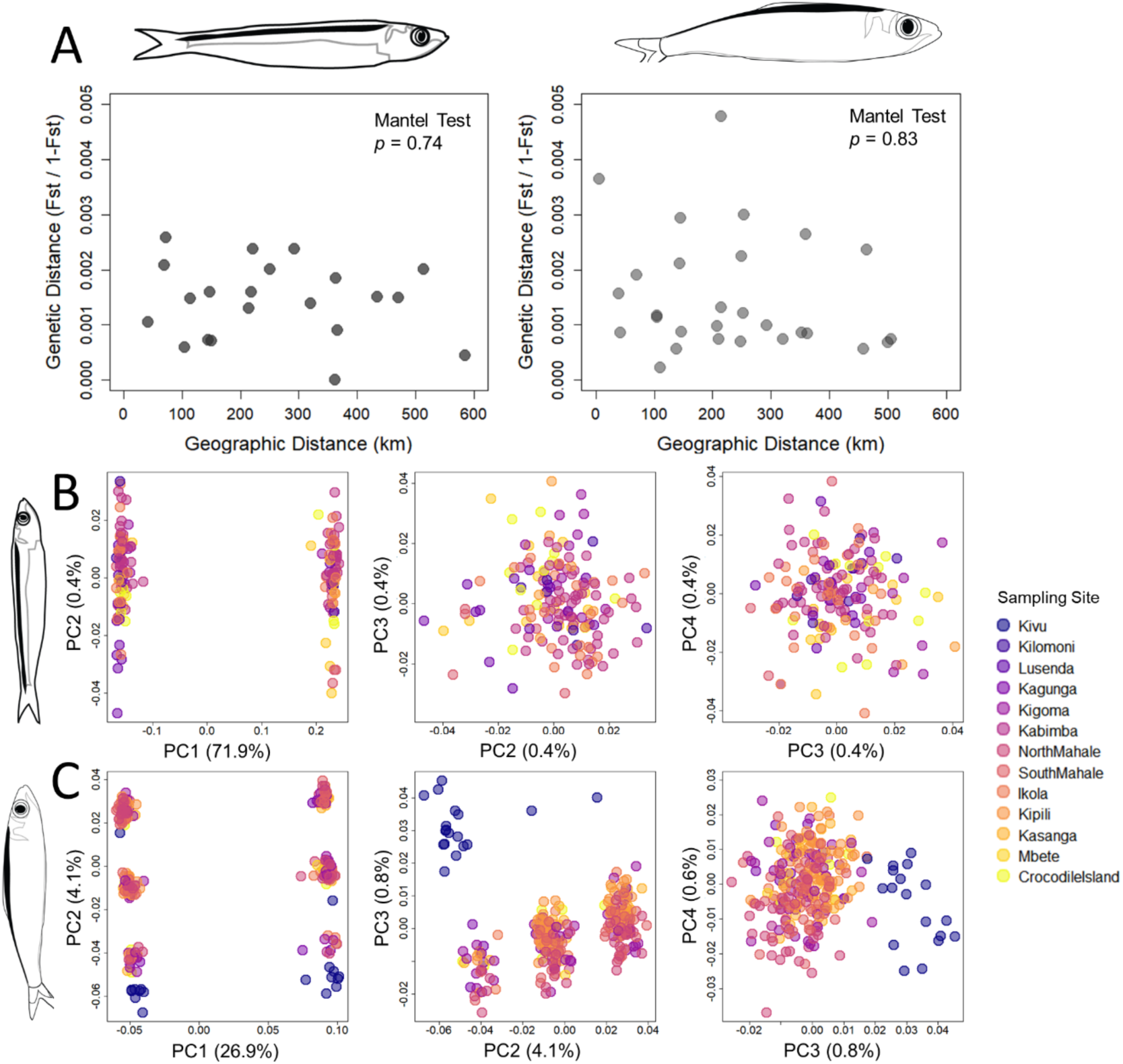
We found no evidence for strong isolation by distance or spatial genetic structure in either species. (A) shows the relationship between genetic and geographic distance for populations *S. tanganicae* (left) and *L. miodon* (right), and the results of Mantel tests between geographic distance (in km) and F_ST_/(1-F_ST_) using the Reich-Patterson F_ST_ estimator. Neither species has evidence for isolation by distance using a Mantel test. (B) and (C) show species-specific principal components analysis of *S. tanganicae* and *L. miodon* individuals, colored by sampling sites. In both species, PC1 differentiates the sexes; in *L. miodon*, PC2 additionally separates each sex into three distinct groups, while PC3 separates individuals from Lake Kivu from those in Lake Tanganyika. Sampling sites are ordered from north (Kivu) to south (Crocodile Island).

**Figure 4.**
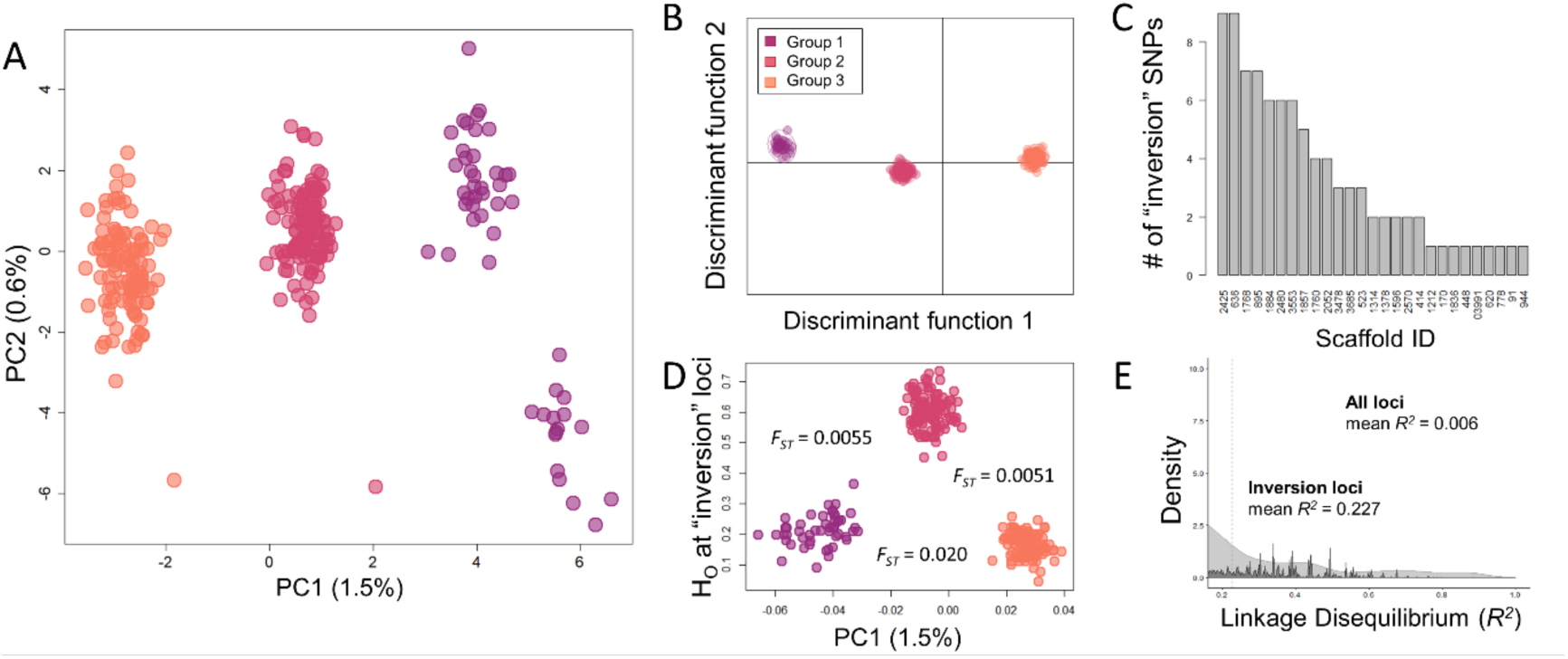
Evidence in *L. miodon* points to the existence of a segregating inversion. (A) Principal component analysis for all *L. miodon* individuals of Lake Tanganyika and Lake Kivu, following the removal of scaffolds containing sex-associated SNPs, demonstrates separation into three groups along the first axis (PC1) and separation between Lake Tanganyika and Lake Kivu individuals along PC2. (B) For discriminant analysis of principal components (DAPC), we assigned individuals according to these three groups and identified SNPs with high loadings along this axis. (C) The 91 SNPs with high loadings along this ‘group’ axis were found on 27 different scaffolds. (D) At these significant loci, two groups were predominantly homozygous, while the third (intermediate) group was generally heterozygous. Divergence values shown between groups were calculated using the Reich-Patterson F_ST_ estimator. Despite the fact that these SNPs were spread out across many scaffolds, the distribution of pairwise linkage disequilibrium values (E) between just the inversion loci (light) has a mean R^2^ = 0.227, whereas the distribution for all loci in the data set for *L. miodon* (dark) has a mean R^2^ = 0.006 (both distributions truncated at R^2^ = 0.2 for better visibility).

With this suggestion of a segregating inversion within *L. miodon*, we tested for Hardy-Weinberg equilibrium among the three groups within Lake Kivu and Lake Tanganyika as a whole and among lake-basins groups (Fig 5). Lake Tanganyika as a whole and each of the basins were in HWE (*X*^2^, p > 0.05). However, the frequencies in Lake Kivu differed significantly from HWE (*X*^2^, p = 0.005) (Fig. 5A and 5B). We additionally found that the proportions of all three karyotype groups differed significantly between Lake Kivu fish and the fish found in each of the north, Mahale (middle), and south basins in Lake Tanganyika (two-proportion z-test; p = 0.010, p = 0.0052, p << 0.001) (Fig. 5B). This result seems to be driven by a much higher frequency of genotype group 3 in Lake Kivu samples than was found in Lake Tanganyika (Fig. 5B). The only significant difference between the three basins within Lake Tanganyika was that the northern basin had a higher frequency of fish with genotype group 3 than either the Mahale or southern basins (two-proportion z-test; p = 0.030, others p > 0.3) (Fig. 5B).

**Figure 5.**
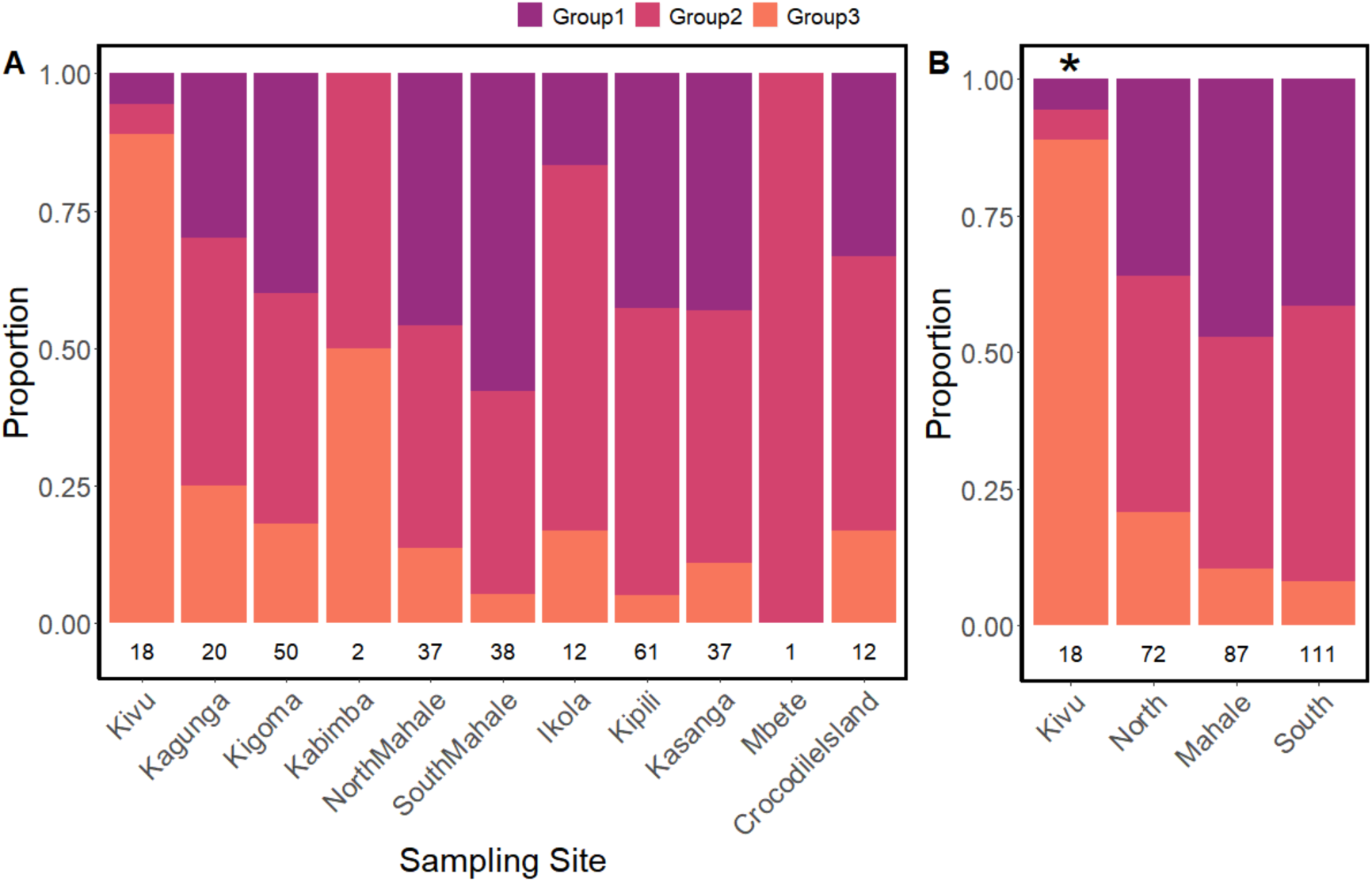
Proportion of individuals in each inversion karyotype group, by sampling site (A) and region (B). Sample sizes of individuals retained in analyses at each sampling site are indicated. In (B), the asterisk (*) indicates rejection of HWE for the region. Sites are ordered by geographic location, from north (Kivu) to south (Crocodile Island). The relative frequency of each haplotype observed differed significantly between Lake Kivu and all three regions in Lake Tanganyika, while only the relative frequency of Group 3 (orange) differed significantly among the three regions within Lake Tanganyika.

### Linkage disequilibrium among identified loci

The distribution of pairwise linkage disequilibrium values among loci in the species-specific data sets were highly right-skewed, with the majority of locus pairs having low to no linkage (overall mean *R*^2^ = 0.007; Fig. 6). In contrast, the subsets of loci identified as sex-linked in the species-specific data sets for *S. tanganicae* and *L. miodon* had mean pairwise LD values of R^2^ = 0.823 and 0.767, respectively (Fig. 6), suggesting that these sets of loci are much more tightly linked than expected based on the distribution of R^2^ values for all loci (Mann-Whitney test; *S. tanganicae* W = 2060242276, p << 0.001; *L. miodon* W = 2954133944, p << 0.001). In *L. miodon*, the inversion group-delineating loci had a mean pairwise LD of R^2^ = 0.227, suggesting that they are also more tightly linked than expected for loci randomly placed in the genome (Mann-Whitney test, W = 682260000000, p << 0.001; Fig 4E), but less tightly linked than the sex-linked loci (Mann-Whitney test, W = 10328070, p << 0.001).

**Figure 6.**
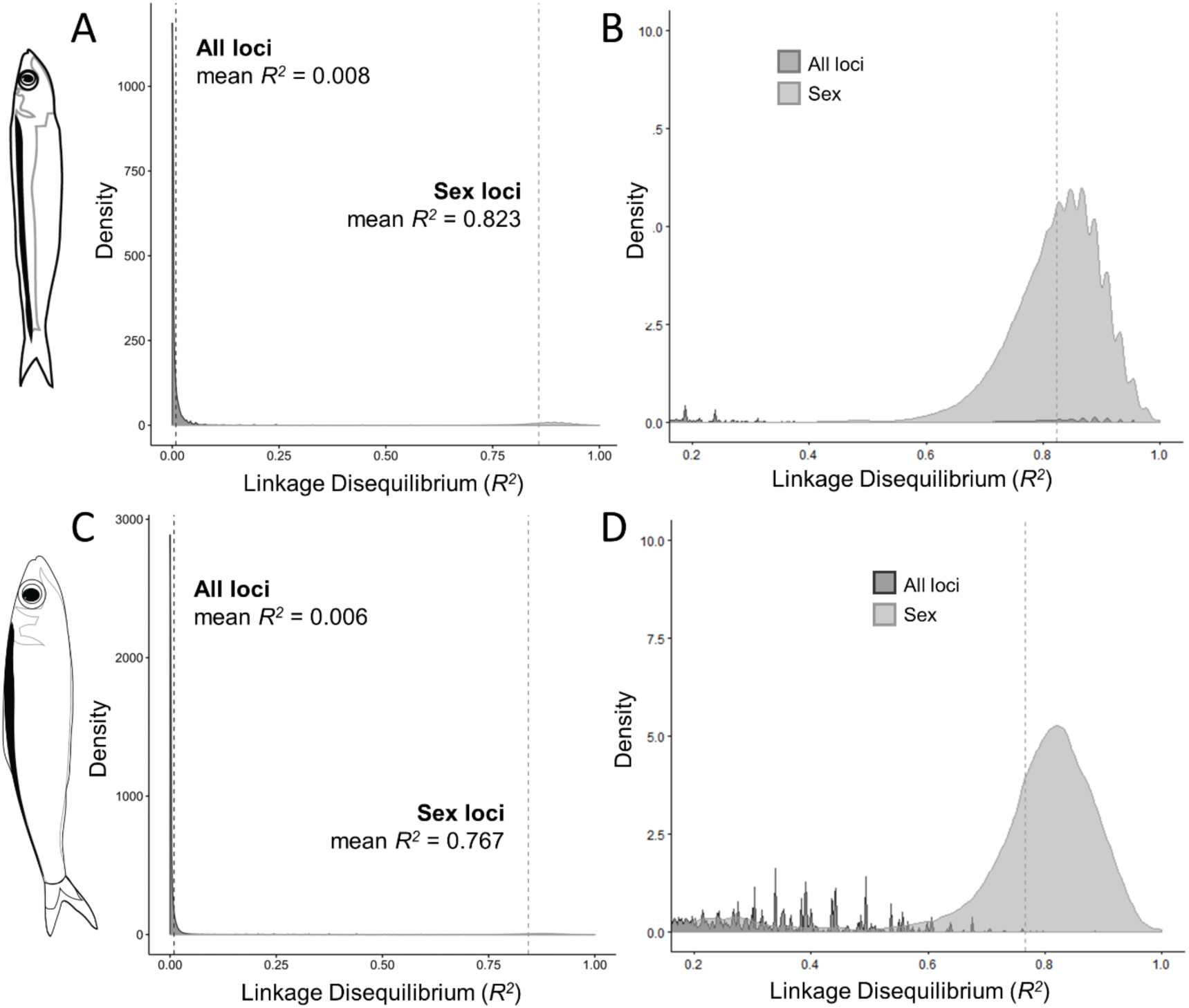
Results from an analysis of pairwise linkage disequilibrium (measured as R^2^) between SNPs in *S. tanganicae* (A-B) and *L. miodon* (C-D), demonstrating that loci associated with sex differences (light gray distributions) are more tightly linked than expected based on linkage values for all loci in the species-specific data sets (dark gray distributions). Panels (B) and (D) have been truncated at R^2^ = 0.2 to better visualize the distribution of LD values for sex-associated SNPs.

### No evidence for isolation-by-distance

Sampling sites throughout the study generally had similar levels of genetic diversity (Θ_W_) for both species (Table 2, Table 3). Within *S. tanganicae*, we found only weak evidence for additional genetic structure beyond the genetic structure linked to sex (Fig. 3B). In contrast, we find very strong genetic structure within each sex in *L. miodon* (Fig. 3C), suggesting the existence of three distinct genetic groups of *L. miodon* in Lake Tanganyika. However, these three groups do not correspond to geographic localities.

**Table 2.** Genetic diversity within (Watterson’s theta, Θ_W_, italicized along diagonal) and differentiation between (Reich-Patterson F_ST_ estimator, above diagonal) sampling sites (unshaded) and basins (shaded) for *S. tanganicae* populations included in this study. Sample sizes are indicated for each sampling location and negative F_ST_ values have been changed to 0. Bolded F_ST_ values are significantly different from 0, based on 100 bootstrapping replicates.

|  | *NORTH* | Kilomoni<br>n=7 | Lusenda<br>n=14 | Kagunga<br>n=12 | Kigoma<br>n=61 | Kabimba<br>n=5 | *MIDDLE* | North<br>Mahale<br>n=25 | South<br>Mahale<br>n=6 | Ikola<br>n=14 | *SOUTH* | Kipili<br>n=12 | Mbete<br>n=17 |
| --- | --- | --- | --- | --- | --- | --- | --- | --- | --- | --- | --- | --- | --- |
| *NORTH* | **0.0044** |  |  |  |  |  | **0.0006** |  |  |  | **0.0006** |  |  |
| Kilomoni |  | *0.0012* | 0.0007 | 0.0003 | 0 | 0.0037 |  | 0.0003 | 0 | 0.0015 |  | 0.0018 | 0.0011 |
| Lusenda |  |  | *0.0007* | **0.0025** | 0.0015 | 0.0023 |  | 0.0015 | 0 | 0.0018 |  | 0.0015 | 0.0004 |
| Kagunga |  |  |  | *0.0007* | 0.0010 | **0.0051** |  | 0.0016 | 0 | 0.0024 |  | 0 | 0.0020 |
| Kigoma |  |  |  |  | *0.0007* | 0.0015 |  | 0.0006 | 0 | **0.0020** |  | 0.0014 | **0.0015** |
| Kabimba |  |  |  |  |  | *0.0008* |  | 0.0018 | 0 | 0.0027 |  | 0.0028 | 0.0009 |
| *MIDDLE* |  |  |  |  |  |  | **0.0014** |  |  |  | **0.0012** |  |  |
| North<br>Mahale |  |  |  |  |  |  |  | *0.0014* | 0 | 0.0007 |  | 0.0013 | 0.0009 |
| South<br>Mahale |  |  |  |  |  |  |  |  | *0.0014* | 0 |  | 0 | 0 |
| Ikola |  |  |  |  |  |  |  |  |  | *0.0010* |  | 0.0021 | **0.0024** |
| *SOUTH* |  |  |  |  |  |  |  |  |  |  | **0.0007** |  |  |
| Kipili |  |  |  |  |  |  |  |  |  |  |  | *0.0011* | 0.00077 |
| Mbete |  |  |  |  |  |  |  |  |  |  |  |  | *0.0011* |

**Table 3.**
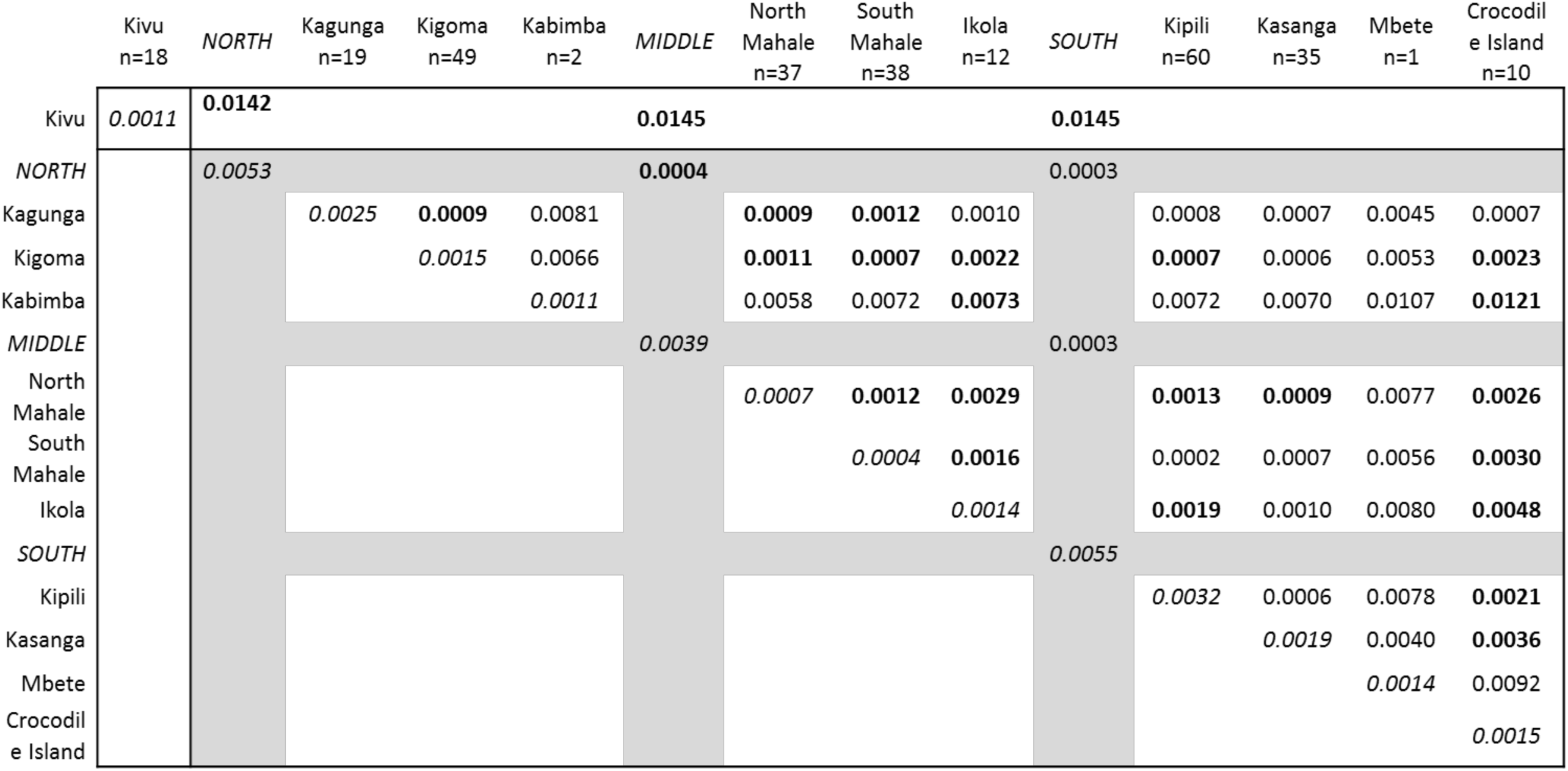
Genetic diversity within (Watterson’s theta, Θ_W_, italicized along diagonal) and differentiation between (Reich-Patterson F_ST_ estimator, above diagonal) sampling sites (unshaded) and basins (shaded) for *L. miodon* populations included in this study. Sample sizes are indicated for each sampling location. Bolded F_ST_ values are significantly different from 0, based on 100 bootstrapping replicates.

In order to examine genetic structure not associated with sex-linked and inversion-linked loci, we removed scaffolds carrying those prior to further analyses. After removing loci associated with sex (in *S. tanganicae*), and those associated with sex and the inversion (in *L. miodon*), 7,235 SNPs remained in the data set for *S. tanganicae* and 17,432 SNPs in the *L. miodon* data set, which were used for spatial structure analyses. We first examined isolation-by-distance patterns with these data sets in both species. While F_ST_ values suggest a weak increase in genetic differentiation with increasing geographic distance (Table 2, Table 3, Fig. 3A), Mantel tests of F_ST_ vs. geographic distance between sampling sites indicated that this association is not significant (*S. tanganicae*, p-value = 0.74; *L. miodon*, p-value = 0.83). The majority of 95% confidence intervals for pairwise F_ST_ estimates overlapped with 0 in *S. tanganicae* and several overlapped 0 in *L. miodon* (Fig. S10). All F_ST_ estimates between basins were small (<0.001), but significantly greater than zero in one out of three comparisons in *L. miodon* and two out of three comparisons in *S. tanganicae* (Fig. S10).

We next conducted analyses of genetic structure using entropy. In both species, the most probable number of genetic groups identified in entropy was K=1 (Fig. S11). In *L. miodon*, the grouping at K=2 separated fish from Lake Kivu from those in Lake Tanganyika. Although we did not detect additional clear clustering in either of the species when using PCA, running entropy at K > 1 for *S. tanganicae* and K > 2 for *L. miodon* identified distinct, albeit not strongly differentiated groups (Fig. S12 and S13). At K = 2 and K = 3 in *S. tanganicae*, multiple genetic groups were present at all sampling sites, but mean group membership for each group differed significantly among sampling sites with more than 2 individuals (one-way ANOVA, F(10,164), all *p* < 0.001; Fig. S14). In *L. miodon*, all groups at K = 3 and K = 4 were present at all sampling sites, except for the Kivu-specific group (Fig. S15). The non-Kivu groups differed significantly in frequency among Tanganyika sampling sites with more than 2 individuals (one-way ANOVA, F(7,242), all p < 0.01; Fig. S14). The Reich F_ST_ estimates for all pairwise comparisons between groups were small but significant in *S. tanganicae* (K=2, mean F_ST_ = 0.0016; K=3 mean F_ST_ = 0.0024; Fig. S16A) and *L. miodon* (K=3 mean non-Kivu F_ST_ = 0.00076; K=4 mean non-Kivu F_ST_ = 0.0012; Fig. S16B).

## Discussion

Little to no spatial genetic structuring is a relatively common observation in pelagic fish species with continuous habitats (e.g. Canales-Aguirre *et al*. 2016; Hutchinson *et al*. 2001; Momigliano *et al*. 2017). However, many studies show that pelagic fish species harbour genetic structure that does not correspond with geographic distance, but instead correlates with ecological adaptation (Berg *et al*. 2017; Kirubakaran *et al*. 2016; Martinez Barrio *et al*. 2016; Pettersson *et al*. 2019) or with cryptic species structure (Doenz *et al*. 2018). We present here the largest genomic data sets analysed for the two freshwater sardines of Lake Tanganyika to date.

We find evidence for the existence of many sex-linked SNPs in both *S. tanganicae* and *L. miodon*, including strong deviations from expected heterozygosity at these loci, suggesting an XY sex determination system with males being the heterogametic sex (Fig. 2). In *L. miodon*, we additionally find three cryptic genetic groups, and patterns in heterozygosity indicate the presence of a segregating chromosomal inversion underlying this genetic structure (Fig. 4 and Fig. S9). All three inversion genotypes (homokaryotypes and heterokaryotype) appear in *L. miodon* from both Lake Tanganyika and Lake Kivu, but relative frequencies of the karyotypes differ among these populations and among regions within Lake Tanganyika (Fig. 5). After removing sex-linked variation and inversion-linked variation, we find no evidence for isolation by distance in *S. tanganicae* or *L. miodon* of Lake Tanganyika, despite the immense size of this lake and extensive geographic sampling of populations of both species. In both species we do find weak genetic structure, with the relative abundance of intraspecific genetic clusters varying between sampling sites.

### Evidence for rapidly evolving genetic XY sex determination in both species

Our results suggest that sex-linked regions of the genome in both *S. tanganicae* and *L. miodon* are large and highly differentiated between males and females (Fig. 2). Despite being located across many scaffolds in our reference genome, these loci are in strong linkage disequilibrium in both species, compared to loci not involved in sex determination (Fig. 6). According to the canonical model of sex chromosome evolution, development of sex chromosomes initiates with the appearance of a sex-determining allele in the vicinity of alternative alleles only favourable for one of the sexes.

Mechanisms reducing recombination, such as inversions that capture the sexually antagonistic locus and the novel sex-determining locus, support the spread of the sex-determining allele in combination with the sexually antagonistic region. Eventually neighboring regions also reduce recombination rate and further mutations accumulate, leading to the formation of a new sex chromosome (Bachtrog 2013; Charlesworth *et al*. 2005; Gammerdinger & Kocher 2018; Wright *et al*. 2016). Examples range from ancient, highly heteromorphic sex chromosomes, to recent neo-sex chromosomes, which are found in mammals (Cortez *et al*. 2014), birds (Graves 2014), and fishes (Feulner *et al*. 2018; Gammerdinger *et al*. 2018; Gammerdinger & Kocher 2018; Kitano & Peichel 2012; Pennell *et al*. 2015; Roberts *et al*. 2009; Ross *et al*. 2009; Yoshida *et al*. 2014). The high number of loci implicated in genetic sex differences in our study, and strong linkage disequilibrium among those loci, in addition to clear patterns of excess heterozygosity in males and homozygosity in females, gives strong indication of the existence of large sex-determining regions in *S. tanganicae* and *L. miodon*, which may lie on sex chromosomes that are distinct for each of the two species.

However, the structural arrangement of these loci remains unclear with our current reference genome. The scaffolds containing sex loci in *L. miodon* total to 76.6Mb, or 13.9% of the reference genome. In contrast, the scaffolds containing sex loci in *S. tanganicae* total 237Mb, which sum to 43% of the reference genome. This suggests that the *S. tanganicae* sex chromosomes are not assembling well to the *L. miodon* genome – rather, they are assembling to many scaffolds to which they do not actually belong. This would be expected if there is little synteny between the sex chromosomes of these species despite their close evolutionary relationship and the nearly equivalent mapping rate of reads from each species to the *L. miodon* genome. It is worth noting that the assembly of sex chromosomes remains challenging due to the haploid nature of sex chromosomes (thus reducing sequencing depth at these regions) and existence of ampliconic and repetitive regions and a high amount of heterochromatin (Tomaszkiewicz *et al*. 2017). Such challenges with assembling sex chromosomes may lead to many scaffolds being implicated in sex determination in initial attempts at assembly, even when these scaffolds do all belong to one chromosome.

Despite large differences in heterozygosity between males and females at sex-associated loci, we did not find systematic differences between males and females in read depth at these loci. If males are the heterogametic sex and the sex chromosomes are strongly differentiated (i.e. the Y chromosome does not assemble to the X chromosome), we would expect males to have half as many reads at sex loci when compared to females. Thus, we expect that some or many Y-chromosome loci are assembling to the X-chromosome in our data set. This further supports the hypothesis that both species likely have young sex chromosomes, where X and Y are not so divergent as to no longer assemble to each other. However, it is difficult to know how much of the Y-chromosome we may be missing in our assembly. In total, we found 502 sex-linked SNPs in *S. tanganicae* and 308 in *L. miodon*. If we relax our SNP filtering thresholds to allow for 80% missing data per site (instead of only 50%), we find an additional 89 SNPs in *S. tanganicae* and 82 SNPs in *L. miodon* that have reads in male fish, but not in females. Some of these sites are located on the same scaffolds already implicated in sex, and others are on scaffolds not yet implicated in sex. These sites likely represent Y chromosome loci that did not assemble well in the reference genome.

We also show that the sets of SNPs linked to sex in *S. tanganicae* and *L. miodon* are entirely distinct from one another, representing strong evidence for rapid evolution in these sex-linked regions (Fig. 2F and S8). Furthermore, since the sex-determining regions in the two species do not appear to be co-located within the same region of the genome, this is evidence that the location of the sex-determining region has shifted. This means that if the common ancestor of these species had a sex-determining region, there appears to have been turnover of sex determining regions in one or both species during the approximately eight million years (95% reliability interval: 2.1–15.9 MYA) since these species diverged (Wilson *et al*. 2008). Rapid turnover of sex chromosomes in closely related species are known from a diversity of taxa (e.g. (Jeffries *et al*. 2018; Kitano & Peichel 2012; Ross *et al*. 2009; Tennessen *et al*. 2018). The proposed mechanisms leading to such rapid turnover rates are chromosomal fusions of an autosome with an already existing sex chromosome, forming a “neo sex chromosome” (Kitano & Peichel 2012; Ross *et al*. 2009) or the translocation of sex loci from one chromosome to another (Tennessen *et al*. 2018). Understanding the mechanisms responsible for the high turnover rate of the sex chromosomes in the Tanganyikan freshwater sardines is a fascinating area for future research.

Furthermore, it will be important for future work to investigate if the strong differentiation between the sexes might also be associated with ecological differences between the sexes. Ecological polymorphism among sexes is known in fishes (Culumber & Tobler 2017; Laporte *et al*. 2018; Parker 1992) and can be ecologically as important as differences between species (Start & De Lisle 2018).

It is worth noting that the strong sex-linked genetic differentiation in *L. miodon* and *S. tanganicae* could have been mistaken for population structure had we filtered our data for excess heterozygosity without first examining it, and had we not been able to carefully phenotypically sex well-preserved, reproductively mature individuals of both species to confirm that the two groups in each species do indeed correspond to sex (Table S1, Fig. 2A). Because of the strong deviations from expected heterozygosity at sex-linked loci, any filtering for heterozygosity would remove these loci from the data set. We believe this may explain why one previous study in *S. tanganicae* using RAD data (De Keyzer *et al*. 2019) did not clearly identify this pattern despite its prevalence in the genome. For organisms with unknown sex determination systems, and for whom sex is not readily identifiable from phenotype, there is danger in conflating biased sampling of the sexes in different populations with population structure in genomic data sets (e.g. Benestan *et al*. 2017). This underscores the importance of sexing sampled individuals whenever possible when analyzing large genomic data sets, and to account for sex in downstream analyses of population or species structure. In our study, the phenotypic and genetic sex were in agreement in all individuals except one *S. tanganicae* individual (Table S1, sample 138863.IKO02). This fish was phenotypically identified as a male but genetically clustered with female individuals, suggesting that this individual was possibly not yet fully mature, and therefore was misidentified phenotypically.

### Evidence for a chromosomal inversion in sympatric L. miodon

Our results reveal the existence of three distinct genetic groups of *L. miodon*. Intriguingly, we find all three of these groups together within the same sampling sites, and even within the same single school of juvenile fishes (Fig. S17). Given patterns of heterozygosity at loci that have high loadings for distinguishing among the genetic clusters (Fig. 4D) together with the strong linkage disequilibrium (Fig. 4E), this structure is consistent with a chromosomal inversion. Chromosomal inversions, first described by Sturtevant (1921), reduce recombination in the inverted region because of the prevention or reduction of crossover in heterogametic individuals (Cooper 1945; Kirkpatrick 2010; Wellenreuther & Bernatchez 2018). Mutations in these chromosomal regions can therefore accumulate independently between the inverted and non-inverted haplotype. Although early work on chromosomal inversions in *Drosophila* has a rich history in evolutionary biology (Kirkpatrick 2010), new genomic sequencing technologies have recently led to inverted regions being detected in many wild species (e.g. Berg *et al*. 2017; Christmas *et al*. 2018; Kirubakaran *et al*. 2016; Lindtke *et al*. 2017; Zinzow-Kramer *et al*. 2015), with implications for the evolution of populations with distinct inversion haplotypes. In *L. miodon*, the strong genetic divergence between the two inversion haplotypes (Fig. 3C, 4A, 4B and 4C) is consistent with this pattern, and indeed the substantial independent evolution of these haplotypes is how the inversion is readily apparent even in a RAD data set. The divergence of the haplotypes, and the high frequency of both of these haplotypes, indicates that this inversion likely did not appear recently, although its apparent absence in *S. tanganicae* indicates it has arisen since the divergence of these sister taxa around 8 million years ago (95% reliability interval: 2.1–15.9 MYA; Wilson *et al*. 2008).

Two issues are of interest given the presence of this chromosomal inversion: first, whether selection was involved in its rise to current frequencies, and second, what evolutionary processes are influencing the current genotype frequencies within populations and the differences in frequencies among populations. Given that both inversion haplotypes appear in relatively high frequencies, it seems unlikely that drift alone could explain the rise of the initially rare derived inversion haplotype to its current frequency in the *L. miodon* population. We expect that *L. miodon* have sustained large effective population sizes through much of their evolutionary history since their split with *S. tanganicae*, implying that drift would have been a continually weak force. Although selection against inversions might occur due to an inversion’s disruption of meiosis or gene expression due to the position of the breakpoints (Kirkpatrick 2010), positive selection may also act on inversions when they carry alleles that themselves are under positive selection. Due to the reduced recombination rates in inversions, these regions of the genome provide opportunities for local or ecological adaptation despite ongoing gene flow or complete random mating (Kirkpatrick & Barton 2006). Although it is unclear given current data whether the inversion that we describe here in *L. miodon* is tied to differential ecological adaptation, this is an important area for future investigation.

Evolutionary pressures on the derived inversion haplotype since its origin may differ from pressures currently acting on either haplotype. When we examine frequencies of the inversion karyotypes pooled across all sampled populations in Lake Tanganyika, the observed frequencies do not differ from Hardy-Weinberg expectations (chi-square = 3.51; p-value = 0.06); this is additionally true within each of the sub-basins within Lake Tanganyika (Fig. 5). However, the Lake Kivu population does show deviation from HWE (chi-square =7.74; p-value = 0.005). Furthermore, frequencies of the inversion karyotypes differ among populations: the proportions of all three karyotypes differ significantly between Lake Kivu and Lake Tanganyika populations, and within Lake Tanganyika, one of the homokaryotypes (Group 3 in Fig. 5), has a higher frequency in the northern basin than in the middle or southern basins (Fig. 5). In Lake Tanganyika, the southern and northern basins differ substantially in nutrient abundance and limnological dynamics, and the Mahale Mountain (middle) region represents the geographical transition between the two basins (Bergamino *et al*. 2010; Kraemer *et al*. 2015; Plisnier *et al*. 1999; Plisnier *et al*. 2009). Thus, it is plausible that differential ecological selection could be driving differences in the frequencies on the inversion karyotypes spatially within the lake, explaining the differences in frequencies we observe among these extant populations. Genetic drift is another possibility to explain the spatial differences in frequencies, and although this is plausible in explaining the frequency differences between Lake Tanganyika and Lake Kivu, it seems a less likely explanation within Lake Tanganyika because of the lack of spatial genetic structure among populations within the lake (Fig. 3A and 3C). Greater understanding of the ecology of these fishes in the north and the south of Lake Tanganyika, and assessment of the genes within the inverted region, is needed to clarify this question.

### Comparing L. miodon populations in Lake Tanganyika to that introduced to Lake Kivu

We found small, but significant divergence between all sub-basin populations of *L. miodon* in Lake Tanganyika compared to the Lake Kivu population (Table 3). In contrast, we found very substantial differences in inversion haplotype frequencies between *L. miodon* in their native Lake Tanganyika and the introduced population in Lake Kivu. This difference in inversion haplotype frequencies could derive from founder effects, from drift within this population since their introduction in the absence of gene flow with the Lake Tanganyika population, or from selection in the Lake Kivu population since their introduction to this substantially different lake environment. We identified individuals in Lake Kivu with all three inversion genotypes that were detected in Lake Tanganyikan fish, suggesting that the inversion is also segregating in Lake Kivu, and that the founding individuals harboured this polymorphism. The difference in haplotype frequencies between Lake Kivu and Lake Tanganyika may have two distinct but not mutually exclusive causes. First, the strong difference in the frequencies of the inversion haplotypes compared to Lake Tanganyika populations may exist due to founder effects. *L. miodon* were introduced to Lake Kivu in the 1950s (Hauser et al. 1995), and all introduced fish were brought from the northern part of Lake Tanganyika. The homokaryotype represented as group 3 in our analyses is the prevalent karyotype in Lake Kivu, and this karyotype also appears in highest frequencies in our samples from northern Lake Tanganyika sites (Fig. 5). Thus, it is plausible that founder effects could have led to the increased frequency of this karyotype within the Lake Kivu population. Second, selection could have contributed to the differences in haplotype frequency between Lake Kivu and Lake Tanganyika. This could either be positive selection for the more common Kivu haplotype, or negative selection against the rarer karyotype caused by low fitness of heterozygotes. For the latter, one possible cause for low heterozygote fitness is disruption in meiosis caused by the inversion, a common scenario for inversions (Kirkpatrick 2010; Kirkpatrick & Barton 2006; Wellenreuther & Bernatchez 2018). However, invoking this mechanism would require understanding why selection due to meiotic dysfunction has not removed the inversion polymorphism in Lake Tanganyika. One possibility would be positive ecological selection on both homokaryotypes in Lake Tanganyika that outweighs selection against heterozygotes due to weak meiotic disfunction; if this ecological selection were released upon introduction to Lake Kivu, selection would then shift entirely to selection against heterozygotes, and would act to decrease the frequency of the less common inversion type. However, this remains speculation and these scenarios need to be examined with additional data.

The current frequency of the inversion karyotypes in Lake Kivu also strongly deviates from Hardy-Weinberg expectations with high frequencies of only one homokaryotype and fewer than expected heterozygotes (Chi-square =7.74; p-value =0.005). This suggests that there are violations to the assumptions of HWE currently operating in the Lake Kivu population. Given that our samples were collected from a geographically proximate location within Lake Kivu and thus are not likely to represent sampling across subpopulations, and given the large population sizes in this lake, we view the most likely violations to be either non-random mating in the Kivu population or selection. As discussed above related to the Lake Kivu haplotype frequencies compared to Lake Tanganyika, the high frequency of the homokaryotype represented as group 3 in our Lake Kivu sample compared to Lake Tanganyika populations may indicate ongoing positive selection for this haplotype or negative selection against the rarer karyotype caused by low fitness of heterozygotes.

In summary, it is likely that the strong shift in genotype frequencies compared to Lake Tanganyika populations and the current deviation from HWE in the Lake Kivu sample is the result of first, founder effects leading to a higher frequency of karyotype 3 in Lake Kivu compared to Lake Tanganyika and second, of selection or non-random mating continuing to distort expected genotype frequencies. Future studies should examine these possibilities more explicitly with a larger sample of individuals from Lake Kivu.

### No isolation-by-distance but weak population structure

For both *L. miodon* and *S. tanganicae*, excluding sex-associated and inversion-associated variation, we find no significant isolation-by-distance when examining the relationship between pairwise genetic differentiation and geographic distance between populations within Lake Tanganyika (Fig. 3a). Pairwise Fsts between sub-basins of Lake Tanganyika are significantly different from zero in two out of three comparisons for *S. tanganicae* and one out of three comparisons for *L. miodon*, however, all values are very small (<0.001; Figure S10). Using entropy to assess genetic structure, analyses for both species support a single genetic group as the most optimal grouping (K=1; Fig. S6). For *L. miodon*, K=2 subdivides the Lake Kivu population from the Lake Tanganyika population. Further examination of results at higher levels of K in both species reveals additional genetic clusters that are composed of sets of individuals (Fig. S12, S13), rather than dividing individuals between clusters, as would be expected if the population were truly panmictic. We interpret this result as implying additional genetic groups in the data which may share allele frequency differences due to selection or drift, or share minor genomic structural variation. Although the groups entropy finds at these values of K do have significant F_ST_s (Fig. S16), their values are small (≤0.015) and the groups are not apparent from PCA, suggesting the effect is due to relatively few small regions in the genome. Because all these groups are distributed across the lake but vary in proportional representation between locations (Fig. S14, S15), it is possible that they differ in allele frequencies due to selection, drift or smaller structural variants.

### Conclusions

Genomic data from *S. tanganicae* and *L. miodon* reveal an interesting array of unexpected patterns in chromosomal evolution. Modern fisheries management seeks to identify locally adapted or otherwise demographically independent units. We do not find significant isolation-by-distance within these two freshwater sardine species from Lake Tanganyika. The strong genetic structure we find is all in sympatry, namely as genetic divergence between the sexes and evidence of a segregating inversion in *L. miodon*. Additionally, we find evidence of weakly differentiated genetic clusters with lake-wide distributions in entropy analyses. From a management perspective, further research should focus on the potential for adaptive differences between the inversion genotypes in *L. miodon* as well as identifying the causes of the additional subtle non-geographical genetic population structure that we found in both species. This study system furthermore offers high potential for further eco-evolutionary study by focusing on the potential for adaptive differences between the sexes in both sardines. All such work will contribute to better understanding the role that these key components of the pelagic community assume in the ecosystem of this lake, which provides important resources to millions of people living at its shores.

## Acknowledgements

This work was funded by the Swiss National Science Foundation (grant CR23I2-166589), a grant from The Nature Conservancy to CEW and PBM, and start-up funding from the University of Wyoming to CEW. CEW was partially supported by NSF grant DEB-1556963. Computing was accomplished with an allocation from the University of Wyoming’s Advanced Research Computing Center, on its Teton Intel x86_64 cluster (https://doi.org/10.15786/M2FY47) and the Genetic Diversity Center (GDC) of ETH Zürich. Special thanks go to Mupape Mukuli for facilitating logistics during fieldwork and to the crew of the MV Maman Benita for fieldwork assistance. We thank the whole team at the Tanzanian Fisheries Research Institute for their support. A special thanks goes to Mary Kishe for her support during fieldwork permission processes and to the Tanzanian Commission for Science and Technology (COSTECH) for their support of this project through permits allowing us to do research in Tanzania. Thanks to Mark Kirkpatrick and his lab group for enlightening discussion regarding the interpretation of these data, to the Wagner lab at the University of Wyoming, and to the FishEc group at EAWAG, especially Kotaro Kagawa and Oliver Selz, for helpful discussion. This manuscript was significantly improved by comments from Erica Larson and three anonymous reviewers.

## Data Accessibility Statement

We will make our genetic data, including our reference genome, publicly available by submitting it to the European Nucleotide Archive (ENA). We intend to submit as soon as possible but by the latest after acceptance of the manuscript. All scripts used for sequence processing and analysis can be found in our Github repository (https://github.com/jessicarick/lake-tanganyika-sardines).

## Author contributions

JJ: developing and writing SNSF grant, sampling and processing fish, identifying phenotypic sex of fish, DNA extractions, preparing RAD libraries, data analysis, writing of the manuscript

JR: sampling and processing fish, DNA extractions, whole genome assembly, data analysis, writing on the manuscript

PBM: developing grant for The Nature Conservancy, contributing samples, discussing results, reviewing manuscript

IK: developing and writing SNSF grant, facilitating permission processes, providing logistics for fieldwork, reviewing manuscript

EAS: sampling and processing fish, facilitating permission processes, providing logistics for fieldwork, enable collaboration with Tanzanian fishermen, discussing manuscript, reviewing manuscript

JBM: sampling and processing fish, facilitating permission processes, providing logistics for fieldwork, enable collaboration with Tanzanian fishermen, discussing and reviewing manuscript

BW: developing and writing SNSF grant, reviewing and discussing manuscript

CD: developing SNSF grant, sampling and processing fish, facilitating logistics during fieldwork, reviewing manuscript

SM: developing SNSF grant, facilitating permission process, RAD library preparation for sequencing, reviewing manuscript

OS: developing and writing SNSF grant, identifying phenotypic sex of fish, facilitating permission process, discussing results, reviewing and discussing manuscript

CEW: developing and writing SNSF and TNC grants, sampling and processing fish, contributing samples, whole genome assembly, data analysis, discussing results, writing manuscript, revise manuscript

## References

Andrews S (2010) FASTQC. A quality control tool for high throughput sequence data.

Bachtrog D (2013) Y-chromosome evolution: emerging insights into processes of Y-chromosome degeneration. Nat Rev Genet 14, 113–124.

Baird NA, Etter PD, Atwood TS, et al. (2008) Rapid SNP discovery and genetic mapping using sequenced RAD markers. PLoS One 3.

Belgrano A, Fowler CW (2011) Ecosystem-based management for Marine Fisheries: An evolving perspective.

Benestan L, Moore JS, Sutherland BJG, et al. (2017) Sex matters in massive parallel sequencing: Evidence for biases in genetic parameter estimation and investigation of sex determination systems. Mol Ecol 26, 6767–6783.

Berg PR, Star B, Pampoulie C, et al. (2017) Trans-oceanic genomic divergence of Atlantic cod ecotypes is associated with large inversions. Heredity (Edinb) 119, 418–428.

Bergamino N, Horion S, Stenuite S, et al. (2010) Spatio-temporal dynamics of phytoplankton and primary production in Lake Tanganyika using a MODIS based bio-optical time series. Remote Sensing of Environment 114, 772–780.

Botsford LW, Castilla JC, Peterson CH (1997) The Management of Fisheries and Marine Ecosystems. Science 277, 509–515.

Canales-Aguirre CB, Ferrada-Fuentes S, Galleguillos R, Hernandez CE (2016) Genetic Structure in a Small Pelagic Fish Coincides with a Marine Protected Area: Seascape Genetics in Patagonian Fjords. PLoS One 11, e0160670.

Chapman DW, van Well P (1978) Growth and Mortality of Stolothrissa tanganicae. Transactions of the American Fisheries Society 107, 26–35.

Charlesworth D, Charlesworth B, Marais G (2005) Steps in the evolution of heteromorphic sex chromosomes. Heredity 95, 118–128.

Catchen J, Hohenlohe PA, Bassham S, Amores A, Cresko WA (2013) Stacks: an analysis tool set for population genomics. Mol Ecol 22, 3124–3140.

Christmas MJ, Wallberg A, Bunikis I, et al. (2018) Chromosomal inversions associated with environmental adaptation in honeybees. Mol Ecol.

Cohen AS, Gergurich EL, Kraemer BM, et al. (2016) Climate warming reduces fish production and benthic habitat in Lake Tanganyika, one of the most biodiverse freshwater ecosystems. Proc Natl Acad Sci U S A 113, 9563–9568.

Cohen AS, Soreghan MJ, Scholz CA (1993) Estimating the age of formation of lakes: an example from Lake Tanganyika, East African Rift system. Geology 21, 511–514.

Collart A (1960) L’introduction du *Stolothrissa tanganicae* (Ndagala) au Lac Kivu. Bulletin Agricole du Congo Belge, 975–985.

Collart A (1989) Introduction et acclimatation de l’isambaza au Lac Kivu. Seminaire “Trente ans apres l’introduction l’isambaza au Lac Kivu “. In: FA0 Report, Gisenyi.

Connallon T, Olito C, Dutoit L, et al. (2018) Local adaptation and the evolution of inversions on sex chromosomes and autosomes. Philos Trans R Soc Lond B Biol Sci 373.

Cooper KW (1945) NORMAL SEGREGATION WITHOUT CHIASMATA IN FEMALE DROSOPHILA MELANOGASTER Genetics 30, 472–484.

Cortez D, Marin R, Toledo-Flores D, et al. (2014) Origins and functional evolution of Y chromosomes across mammals. Nature 508, 488.

Coulter GW (1976) The biology of Lates species (Nile perch) in Lake Tanganyika, and the status of the pelagic fishery for Lates species and *Luciolates stappersii* (Blgr.). Journal of Fish Biology 9, 235–259.

Coulter GW (1991) Lake Tanganyika and its Life British Museum (Natural History) Cromwell Road, London SW7 5BD & Oxford University Press, Walton Street, Oxford OX2 6DP.

Culumber ZW, Tobler M (2017) Sex-specific evolution during the diversification of live-bearing fishes. Nature Ecology & Evolution 1, 1185–1191.

Danecek P, Auton A, Abecasis G, et al. (2011) The variant call format and VCFtools. Bioinformatics 27, 2156–2158.

De Keyzer ELR, De Corte Z, Van Steenberge M, et al. (2019) First genomic study on Lake Tanganyika sprat *Stolothrissa tanganicae*: a lack of population structure calls for integrated management of this important fisheries target species. BMC Evol Biol 19, 6.

Doenz CJ, Bittner D, Vonlanthen P, Wagner CE, Seehausen O (2018) Rapid buildup of sympatric species diversity in Alpine whitefish. Ecology and Evolution 8, 9398–9412.

Ellis CMA (1971) The size at maturity and breeding seasons of sardines in southern Lake Tanganyika. African Journal of Tropical Hydrobiology and Fisheries 1, 59–66.

FAO (1995) Management of African inland fisheries for sustainable production. In: First Pan African Fisheries Congress and Exhibition. FAO Rome, UNEP, NAIROBI.

Feulner PGD, Schwarzer J, Haesler MP, Meier JI, Seehausen O (2018) A Dense Linkage Map of Lake Victoria Cichlids Improved the Pundamilia Genome Assembly and Revealed a Major QTL for Sex-Determination. G3 (Bethesda) 8, 2411–2420.

Gammerdinger WJ, Conte MA, Sandkam BA, et al. (2018) Novel Sex Chromosomes in 3 Cichlid Fishes from Lake Tanganyika. J Hered 109, 489–500.

Gammerdinger WJ, Kocher TD (2018) Unusual Diversity of Sex Chromosomes in African Cichlid Fishes. Genes (Basel) 9.

Graves JAM (2014) Avian sex, sex chromosomes, and dosage compensation in the age of genomics. Chromosome Research 22, 45–57.

Hauser L, Carvalho GR, Pitcher TJ (1995) Morphological and genetic differentiation of the African clupeid Limnothrissa miodon 34 years after introduction to Lake Kivu. Journal of Fish Biology 47, 127 144.

Hauser L, Carvalho GR, Pitcher TJ (1998) Genetic population structure in the Lake Tanganyika sardine *Limnothrissa miodon*. Journal of Fish Biology 53, 413–429.

Hooper DM, Price TD (2015) Rates of karyotypic evolution in Estrildid finches differ between island and continental clades. Evolution 69, 890–903.

Hooper DM, Griffith SC, Price TD (2019) Sex chromosome inversions enforce reproductive isolation across an avian hybrid zone. Mol Ecol 28, 1246–1262.

Hutchinson WF (2008) The dangers of ignoring stock complexity in fishery management: the case of the North Sea cod. Biol Lett 4, 693–695.

Hutchinson WF, Carvalho GR, Rogers SI (2001) Marked genetic structuring in localised spawning populations of cod *Gadus morhua* in the North Sea and adjoining waters, as revealed by microsatellites. Marine Ecology Progress Series 223, 251–269.

Jeffries DL, Lavanchy G, Sermier R, et al. (2018) A rapid rate of sex-chromosome turnover and non-random transitions in true frogs. Nat Commun 9, 4088.

Jombart T (2008) adegenet: a R package for the multivariate analysis of genetic markers. Bioinformatics 24, 1403–1405.

Jombart T, Devillard S, Balloux F (2010) Discriminant analysis of principal components: a new method for the analysis of genetically structured populations. BMC Genetics 11, 94.

Jones FC, Grabherr MG, Chan YF, et al. (2012) The genomic basis of adaptive evolution in threespine sticklebacks. Nature 484, 55–61.

Kimirei IA, Mgaya YD, Chande AI (2008) Changes in species composition and abundance of commercially important pelagic fish species in Kigoma area, Lake Tanganyika, Tanzania. Aquatic Ecosystem Health and Management 11, 29–35.

Kirkpatrick M (2010) How and Why Chromosome Inversions Evolve. PLoS Biology 8, e1000501.

Kirkpatrick M, Barton N (2006) Chromosome inversions, local adaptation and speciation. Genetics 173, 419–434.

Kirubakaran TG, Andersen Ø, De Rosa MC, et al. (2019) Characterization of a male specific region containing a candidate sex determining gene in Atlantic cod. Scientific Reports 9, 116–116.

Kirubakaran TG, Grove H, Kent MP, et al. (2016) Two adjacent inversions maintain genomic differentiation between migratory and stationary ecotypes of Atlantic cod. Mol Ecol 25, 2130–2143.

Kitano J, Peichel CL (2012) Turnover of sex chromosomes and speciation in fishes. Environ Biol Fishes 94, 549–558.

Korneliussen TS, Albrechtsen A, Nielsen R (2014) ANGSD: Analysis of Next Generation Sequencing Data. BMC Bioinformatics 15, 356.

Korneliussen TS, Moltke I, Albrechtsen A, Nielsen R (2013) Calculation of Tajima’s D and other neutrality test statistics from low depth next-generation sequencing data. BMC Bioinformatics 14, 289.

Kraemer BM, Hook S, Huttula T, et al. (2015) Century-Long Warming Trends in the Upper Water Column of Lake Tanganyika. PLoS One 10, e0132490.

Kurki H, Mannini P, Vuorinen I, et al. (1999) Macrozooplankton communities in Lake Tanganyika indicate food chain differences between the northern part and the main basins. Hydrobiologia 407, 123–129.

Kuusipalo L (1999) Genetic variation in the populations of pelagic clupeids Stolothrissa tanganicae and Limnothrissa miodon and nile perch (Lates stappersii, L. mariae) in Lake Tanganyika. FAO/FINNIDA Research for the Management of the Fisheries of Lake Tanganyika., p. 28p.

Lamichhaney S, Fuentes-Pardo AP, Rafati N, et al. (2017) Parallel adaptive evolution of geographically distant herring populations on both sides of the North Atlantic Ocean. Proceedings of the National Academy of Sciences 114, E3452–E3461.

Langmead B, Salzberg SL (2012) Fast gapped-read alignment with Bowtie 2. Nat Methods 9, 357–359.

Laporte M, Berrebi P, Claude J, et al. (2018) The ecology of sexual dimorphism in size and shape of the freshwater blenny *Salaria fluviatilis*. Curr Zool 64, 183–191.

Li H, Durbin R (2009) Fast and accurate short read alignment with Burrows–Wheeler transform. Bioinformatics 25, 1754–1760.

Li H, Handsaker B, Wysoker A, et al. (2009) The Sequence Alignment/Map format and SAMtools. Bioinformatics 25, 2078–2079.

Lindtke D, Lucek K, Soria-Carrasco V, et al. (2017) Long-term balancing selection on chromosomal variants associated with crypsis in a stick insect. Mol Ecol 26, 6189–6205.

Loiselle S, Cózar A, Adgo E, et al. (2014) Decadal Trends and Common Dynamics of the Bio-Optical and Thermal Characteristics of the African Great Lakes. PLoS One 9, e93656.

Mannini P, Aro E, Katonda I, et al. (1996) Pelagic fish stocks of Lake Tanganyika: biology and exploitation. FAO/FINNIDA Research for the Management of the Fisheries of Lake Tanganyika. In: GCP/RAF/271/FIN—TD/53 (En), p. 60p.

Marques DA, Lucek K, Meier JI, et al. (2016) Genomics of Rapid Incipient Speciation in Sympatric Threespine Stickleback. PLoS Genet 12, e1005887.

Martinez Barrio A, Lamichhaney S, Fan G, et al. (2016) The genetic basis for ecological adaptation of the Atlantic herring revealed by genome sequencing. Elife 5.

Martinez PA, Zurano JP, Amado TF, et al. (2015) Chromosomal diversity in tropical reef fishes is related to body size and depth range. Molecular Phylogenetics and Evolution 93, 1–4.

McGlue MM, Lezzar KE, Cohen AS, et al. (2007) Seismic records of late Pleistocene aridity in Lake Tanganyika, tropical East Africa. Journal of Paleolimnology 40, 635–653.

Mikheenko A, Prjibelski A, Saveliev V, Antipov D, Gurevich A (2018) Versatile genome assembly evaluation with QUAST-LG. Bioinformatics 34, i142–i150.

Mölsä H, Reynolds JE, Coenen EJ, Lindqvist OV (1999) Fisheries research towards resource management on Lake Tanganyika. Hydrobiologia 407, 1–24.

Mölsä H, Sarvala J, Badende S, et al. (2002) Ecosystem monitoring in the development of sustainable fisheries in Lake Tanganyika. Aquatic Ecosystem Health and Management 5, 267–281.

Momigliano P, Jokinen H, Fraimout A, et al. (2017) Extraordinarily rapid speciation in a marine fish. Proceedings of the National Academy of Sciences 114, 6074–6079.

Mulimbwa N, Raeymaekers JAM, Sarvala J (2014a) Seasonal changes in the pelagic catch of two clupeid zooplanktivores in relation to the abundance of copepod zooplankton in the northern end of Lake Tanganyika. Aquatic Ecosystem Health and Management 17, 25–33.

Mulimbwa N, Sarvala J, Raeymaekers JAM (2014b) Reproducitve activities of two zooplanktivorous clupeid fish in relation to the seasonal abundance of copepod prey in the northern end of Lake Tanganyika. Belgian Journal of Zoology 144, 77–92.

Natri HM, Merila J, Shikano T (2019) The evolution of sex determination associated with a chromosomal inversion. Nat Commun 10, 145.

O’Reilly CM, Alin SR, Piisnier PD, Cohen AS, McKee BA (2003) Climate change decreases aquatic ecosystem productivity of Lake Tanganyika, Africa. Nature 424, 766–768.

Parker GA (1992) The evolution of sexual size dimorphism in fish. Journal of Fish Biology, 1–20.

Pearce MJ (1985) A description and stock assessment of the pelagic fishery in the South-east arm of the Zambian waters of Lake Tanganyika. In: Report of the department of fisheries, Zambian pp. 1–74.

Pennell MW, Kirkpatrick M, Otto SP, et al. (2015) Y Fuse? Sex Chromosome Fusions in Fishes and Reptiles. PLOS Genetics 11, e1005237.

Pettersson ME, Rochus CM, Han F, et al. (2019) A chromosome-level assembly of the Atlantic herring genome-detection of a supergene and other signals of selection. Genome Research 29, 1919–1928.

Plisnier PD, Chitamwebwa D, Mwape L, et al. (1999) Limnological annual cycle inferred from physical-chemical fluctuations at three stations of Lake Tanganyika. Hydrobiologia 407, 45–58.

Plisnier PD, Mgana H, Kimirei I, et al. (2009) Limnological variability and pelagic fish abundance (*Stolothrissa tanganicae* and *Lates stappersii*) in Lake Tanganyika. Hydrobiologia 625, 117–134.

Presgraves DC (2008) Sex chromosomes and speciation in Drosophila. Trends in Genetics 24, 336–343.

Pritchard JK, Przeworski M (2001) Linkage Disequilibrium in Humans: Models and Data. The American Journal of Human Genetics 69, 1–14.

Purcell S, Neale B, Todd-Brown K, et al. (2007) PLINK: A Tool Set for Whole-Genome Association and Population-Based Linkage Analyses. The American Journal of Human Genetics 81, 559–575.

Qvarnstrom A, Bailey RI (2009) Speciation through evolution of sex-linked genes. Heredity (Edinb) 102, 4–15.

Reich D, Thangaraj K, Patterson N, Price AL, Singh L (2009) Reconstructing Indian population history. Nature 461, 489–494.

Roberts RB, Ser JR, Kocher TD (2009) Sexual Conflict Resolved by Invasion of a Novel Sex Determiner in Lake Malawi Cichlid Fishes. Science 326, 998–1001.

Roesti M, Kueng B, Moser D, Berner D (2015) The genomics of ecological vicariance in threespine stickleback fish. Nat Commun 6, 8767.

Ross JA, Urton JR, Boland J, Shapiro MD, Peichel CL (2009) Turnover of Sex Chromosomes in the Stickleback Fishes (Gasterosteidae). PLOS Genetics 5, e1000391.

Sarvala J, Tarvainen M, Salonen K, Mölsä H (2002) Pelagic food web as the basis of fisheries in Lake Tanganyika: A bioenergetic modeling analysis. Aquatic Ecosystem Health and Management 5, 283–292.

Spliethoff PC, de Longh HH, Frank VG (1983) Success of the Introduction of the Fresh Water Clupeid *Limnothrissa miodon* (Boulenger) in Lake Kivu. Aquaculture Research 14, 17–31.

Start D, De Lisle S (2018) Sexual dimorphism in a top predator (*Notophthalmus viridescens*) drives aquatic prey community assembly. Proc Biol Sci 285.

Sturtevant AH (1921) A Case of Rearrangement of Genes in Drosophila. Genetics 7, 235–237.

Tennessen JA, Wei N, Straub SCK, et al. (2018) Repeated translocation of a gene cassette drives sex-chromosome turnover in strawberries. PLoS Biol 16, e2006062.

Tomaszkiewicz M, Medvedev P, Makova KD (2017) Y and W Chromosome Assemblies: Approaches and Discoveries. Trends Genet 33, 266–282.

van der Knaap M (2013) Comparative analysis of fisheries restoration and public participation in Lake Victoria and Lake Tanganyika. Aquatic Ecosystem Health and Management 16, 279–287.

Van der Knaap M, Katonda KI, De Graaf GJ (2014) Lake Tanganyika fisheries frame survey analysis: Assessment of the options for management of the fisheries of Lake Tanganyika. Aquatic Ecosystem Health and Management 17, 4–13.

van Zwieten PAM, Roest FC, Machiels MAM, Van Densen WLT (2002) Effects of inter-annual variability, seasonality and persistence on the perception of long-term trends in catch rates of the industrial pelagic purse-seine fishery of northern Lake Tanganyika (Burundi). Fisheries Research 54, 329–348.

Verburg P, Antenucci JP, Hecky RE (2011) Differential cooling drives large-scale convective circulation in Lake Tanganyika. Limnology and Oceanography 56, 910–926.

Verburg P, Hecky RE (2003) Wind patterns, evaporation, and related physical variables in Lake Tanganyika, east Africa. Journal of Great Lakes Research 29, 48–61.

Verburg P, Hecky RE, Kling H (2003) Ecological consequences of a century of warming in Lake Tanganyika. Science 301, 505–507.

Wellenreuther M, Bernatchez L (2018) Eco-Evolutionary Genomics of Chromosomal Inversions. Trends Ecol Evol 33, 427–440.

Williams JGK, Kubelik AR, Livak KJ, Rafalski JA, Tingey SV (1990) DNA polymorphisms amplified by arbitrary primers are useful as genetic markers. Nucleic Acids Research 18, 6531–6535.

Wilson AB, Teugels GG, Meyer A (2008) Marine incursion: the freshwater herring of Lake Tanganyika are the product of a marine invasion into West Africa. PLoS One 3, e1979.

Wright AE, Dean R, Zimmer F, Mank JE (2016) How to make a sex chromosome. Nature Communications 7, 12087.

Yoshida K, Makino T, Yamaguchi K, et al. (2014) Sex Chromosome Turnover Contributes to Genomic Divergence between Incipient Stickleback Species. PLOS Genetics 10, e1004223.

Zheng X, Levine D, Shen J, et al. (2012) A high-performance computing toolset for relatedness and principal component analysis of SNP data. Bioinformatics 28, 3326–3328.

Zinzow-Kramer WM, Horton BM, McKee CD, et al. (2015) Genes located in a chromosomal inversion are correlated with territorial song in white-throated sparrows. Genes Brain Behav 14, 641–654.

