## Supplemental Tables and Figures for "Structural genomic variation leads to unexpected genetic differentiation in Lake Tanganyika’s sardines"

**Supplemental Table**

**Table S1.** Morphological sexing results in *S. tanganicae* and *L. miodon*. Mature formalin-preserved individuals were dissected and visually inspected for the presence of gonads to determine phenotypic sex. SL = standard length.

| ***S. tanganicae*** |  |  |  | ***L. miodon*** |  |  |  |
| --- | --- | --- | --- | --- | --- | --- | --- |
| **Sample ID** | **Genetic sex** | **Phenotypic sex** | **SL (mm)** | **Sample ID** | **Genetic sex** | **Phenotypic sex** | **SL (mm)** |
| 138863.IKO02 | female | male | 99 | 139100.KAS26 | male | male | 105 |
| 138864.IKO03 | male | male | 99 | 139122.KAS48 | female | female | 105 |
| 138866.IKO05 | female | female | 100 | 138826.KAT25 | female | female | 91 |
| 138867.IKO06 | female | female | 100 | 138827.KAT26 | female | female | 99 |
| 138868.IKO07 | male | male | 102 | 138828.KAT27 | female | female | 98 |
| 138869.IKO08 | male | male | 94 | 138831.KAT30 | female | female | 87 |
| 138870.IKO09 | female | female | 101 | 138832.KAT31 | female | female | 86 |
| 138872.IKO11 | male | male | 99 | 138836.KAT35 | female | female | 84 |
| 138873.IKO12 | female | female | 99 | 138842.KAT41 | female | female | 90 |
| 138874.IKO13 | female | female | 99 | 138919.IKO58 | male | male | 130 |
| 138883.IKO22 | male | male | 98 | 138955.IKO94 | male | male | 104 |
| 138889.IKO28 | male | male | 101 | 138982.KIP15 | female | female | 102 |
| 139217.KAG51 | female | female | 75 | 138994.KIP26 | female | female | 109 |
| 139219.KAG53 | female | female | 72 | 138998.KIP30 | female | female | 92 |
|  |  |  |  | 139010.KIP42 | female | female | 94 |
|  |  |  |  | 139011.KIP43 | female | female | 102 |
|  |  |  |  | 139022.KIP54 | female | female | 99 |
|  |  |  |  | 139098.KAS24 | male | male | 105 |
|  |  |  |  | 139101.KAS27 | female | female | 108 |
|  |  |  |  | 139117.KAS43 | female | female | 114 |
|  |  |  |  | 139119.KAS45 | male | male | 105 |
|  |  |  |  | 139137.KAS56 | male | male | 106 |
|  |  |  |  | 139243.KAG77 | male | male | 91 |
|  |  |  |  | 139245.KAG79 | female | female | 102 |
|  |  |  |  | 139246.KAG80 | male | male | 100 |
|  |  |  |  | 139252.KAG86 | male | male | 90 |
|  |  |  |  | 64310.KIV01 | female | female | 123 |
|  |  |  |  | 64311.KIV02 | female | female | 119 |
|  |  |  |  | 64312.KIV03 | female | female | 108 |
|  |  |  |  | 64445.KIV04 | male | male | 93 |
|  |  |  |  | 64450.KIV05 | male | male | 86 |
|  |  |  |  | 64452.KIV06 | male | male | 87 |
|  |  |  |  | 64554.KIV07 | male | male | 80 |
|  |  |  |  | 64555.KIV08 | male | male | 75 |
|  |  |  |  | 64556.KIV09 | male | male | 76 |
|  |  |  |  | 64557.KIV10 | female | female | 86 |
|  |  |  |  | 64558.KIV11 | male | male | 82 |
|  |  |  |  | 64559.KIV12 | male | male | 86 |
|  |  |  |  | 64593.KIV13 | female | female | 105 |
|  |  |  |  | 64595.KIV14 | female | female | 74 |
|  |  |  |  | 64596.KIV15 | female | female | 75 |
|  |  |  |  | 64597.KIV16 | female | female | 77 |
|  |  |  |  | 64598.KIV17 | female | female | 80 |
|  |  |  |  | 64599.KIV18 | female | female | 75 |
|  |  |  |  | 64761.KIV19 | male | male | 74 |

**Supplemental Figures**

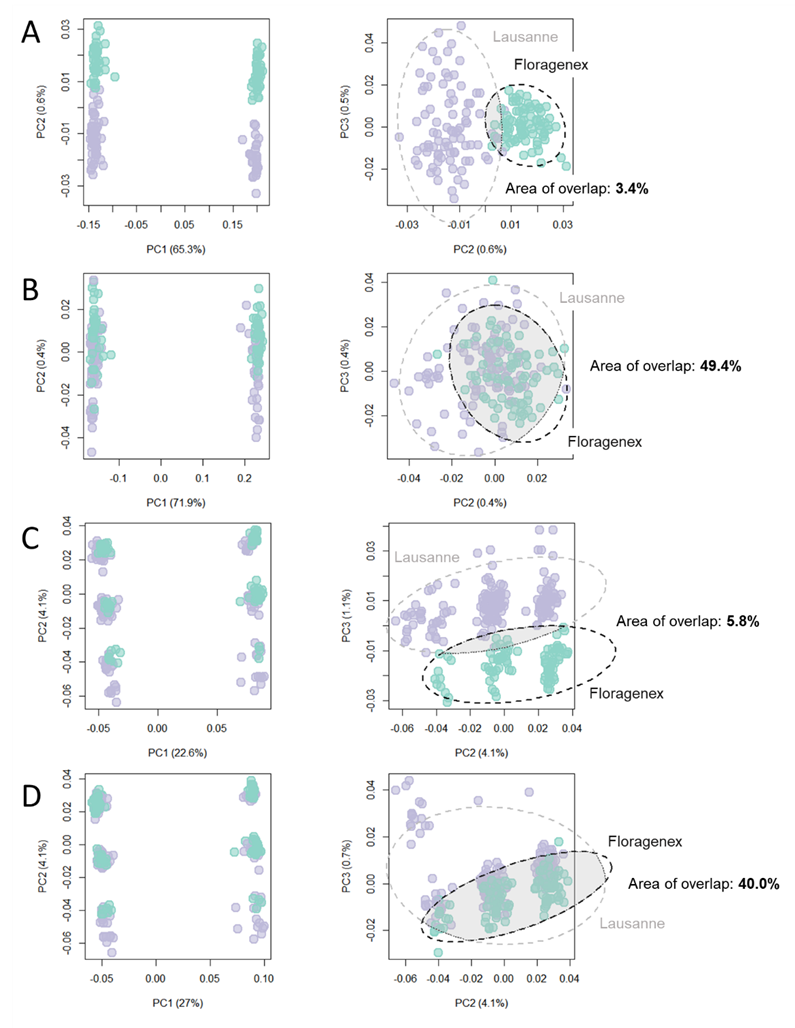

**Figure S1**. Principal component analysis of genetic data for *S. tanganicae* and *L. miodon* individuals, colored by sequencing facility. (A) and (C) are original data sets, prior to filtering for library effects in *S. tanganicae* and *L. miodon*, respectively. (B) and (D) show results after removing SNPs occurring in only one group of samples or the other, demonstrating the removal of library effects. Dashed lines show 95% confidence ellipses for the samples from each sequencing facility, with the overlap between these ellipses shaded in gray to demonstrate an improvement in overlap between the unfiltered and filtered data sets. We do not expect complete overlap, even if all library effects are removed, due to individual differences in the samples. In both species, major features of the data (i.e. sex on PC1, three groups on PC2 in *L. miodon*) remain unchanged following filtering.

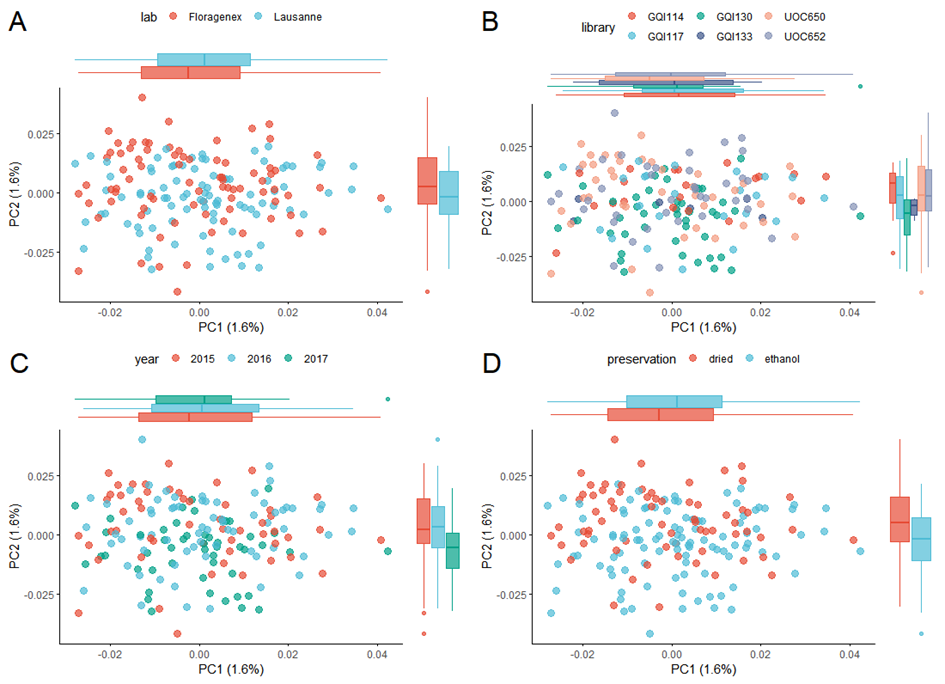

**Figure S2**. Genetic PCA of *Stolothrissa tanganicae* individuals, colored by (A) sequencing facility; (B) RAD library; (C) year collected; and (D) tissue preservation method, demonstrating that these differences in sample treatment are not reflected in population structure results. Boxplots on the margins of each plot show the distribution of PC1 and PC2 values for each group. All PCAs were performed using the SNP data set omitting scaffolds containing sex-linked loci and corrected for sequencing effects.

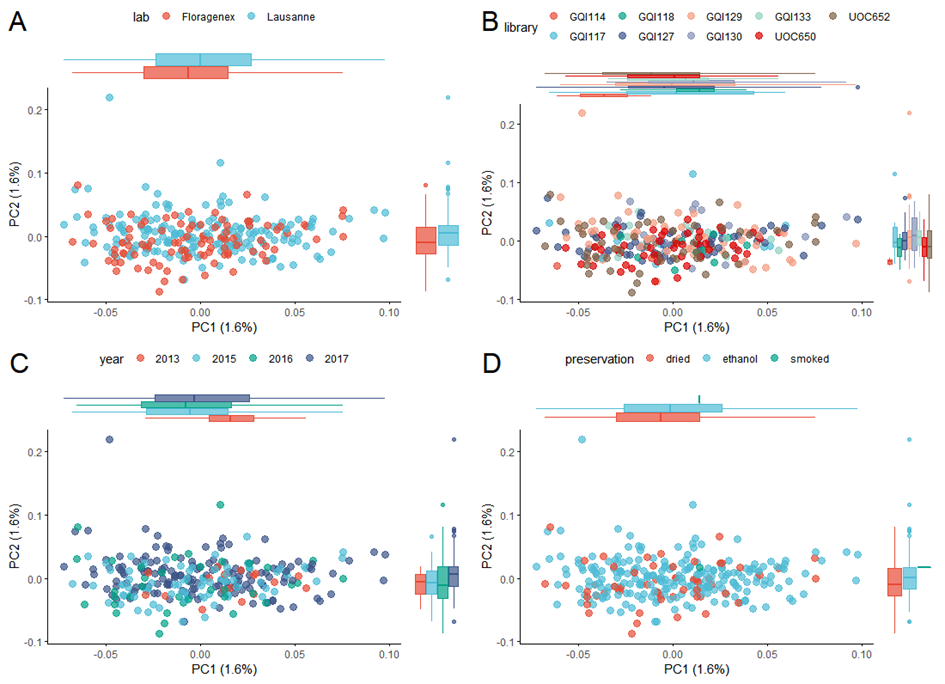

**Figure S3**. Genetic PCA of *Limnothrissa miodon* individuals, colored by (A) sequencing facility; (B) RAD library; (C) year collected; and (D) tissue preservation method, demonstrating that these differences in sample treatment are not reflected in population structure results. Boxplots on the margins of each plot show the distribution of PC1 and PC2 values for each group. All PCAs were performed using the SNP data set omitting scaffolds containing inversion- and sex-linked loci and corrected for sequencing effects.

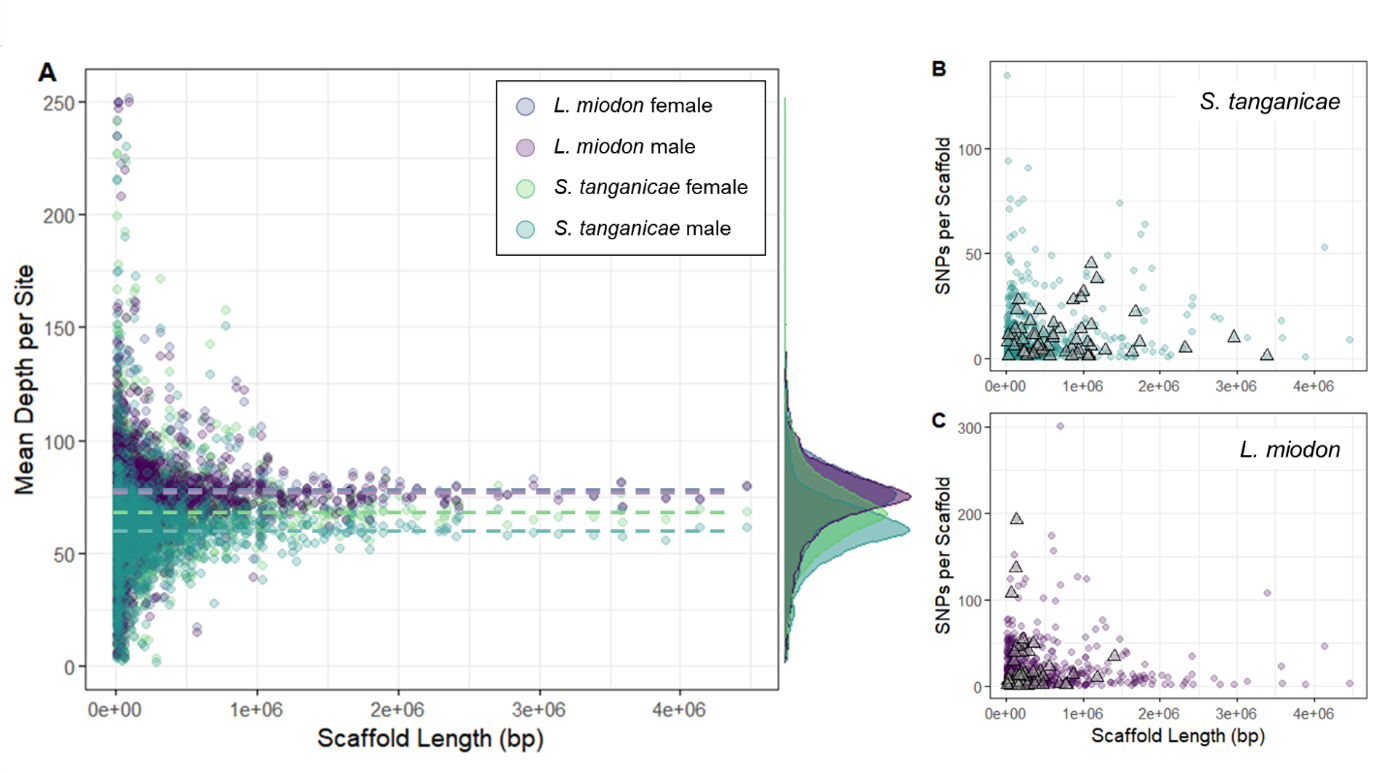

**Figure S4**. Coverage and number of SNPs found for each scaffold, for each species. (A) The mean number of reads per site across a scaffold did not differ significantly with scaffold length in either species, demonstrating good coverage of scaffolds, particularly in long scaffolds. Coverage did vary among species and among sexes within species. *L. miodon* females and males generally had similar read depths (female mean = 73 reads per site; male mean = 73), followed by *S. tanganicae* females (mean = 68 reads per site), and *S. tanganicae* males (mean = 61 reads per site). However, this did not translate into large differences in the number of SNPs per scaffold when comparing (B) *S. tanganicae* to (C) *L. miodon*. In (B) and (C), triangles indicate scaffolds containing sex SNPs in the indicated species.

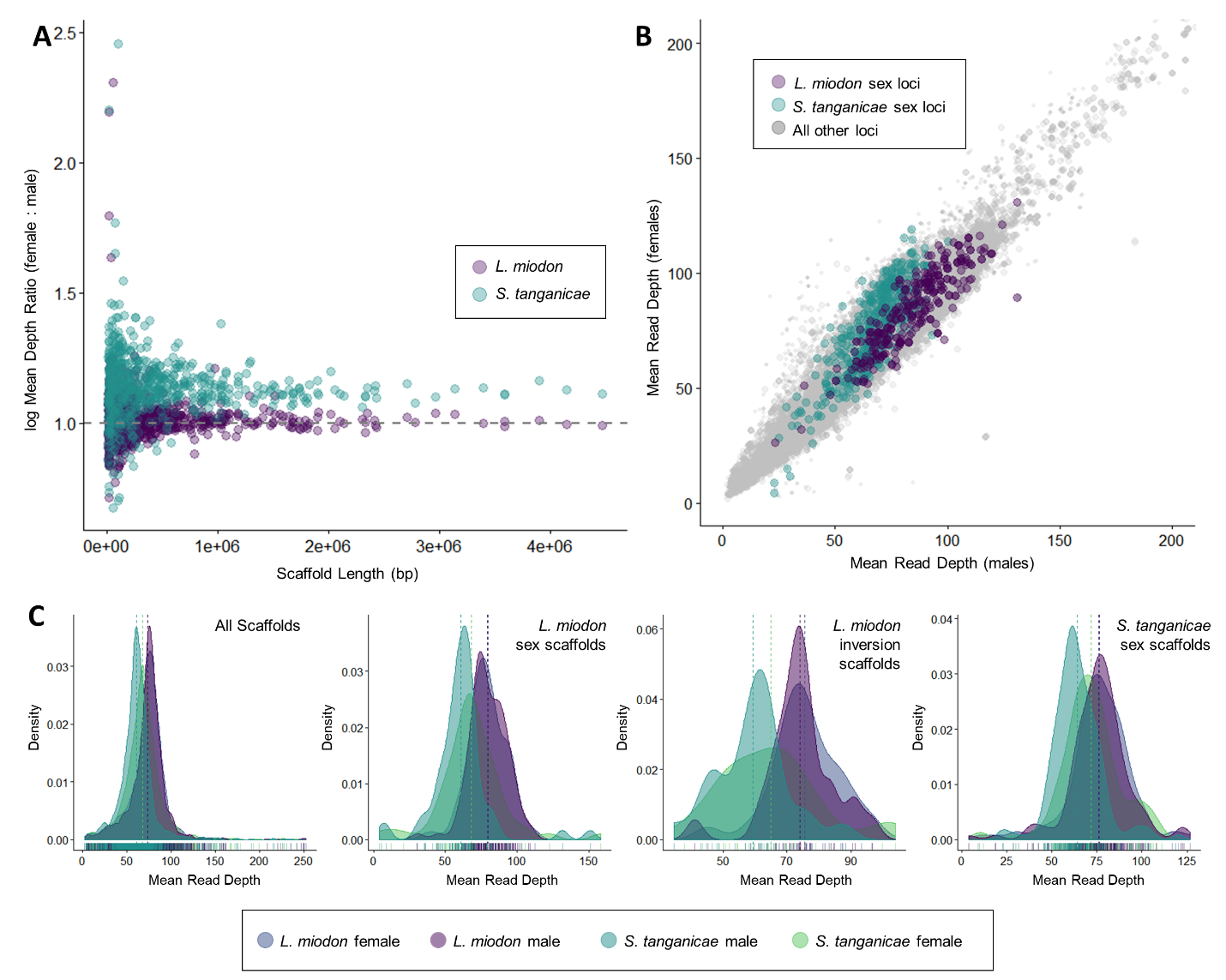

**Figure S5**. In both species, mean read depth is similar between scaffolds carrying sex linked or inversion linked SNPs and scaffolds carrying none of these. Plots show (A) the ratio of mean depth per scaffold between female and male individuals for *L. miodon* (purple) and *S. tanganicae* (teal), for all scaffolds. Dashed line indicates a 1:1 (female:male) ratio of coverage. (B) Relationship between mean read depth at individual SNPs in males and mean read depth in females, for all loci (gray), *L. miodon* sex loci in *L. miodon* (purple), and *S. tanganicae* sex loci in *S. tanganicae* (teal). The ratios between mean depth for males and females did not differ at sex loci in either species when compared to the ratios for all loci. Panel (C) shows the distribution of mean read depths for each scaffold (after removing sites with read depth < 2), using (from left to right) the set of all scaffolds (n = 5,515 scaffolds), scaffolds containing SNPs linked to sex in *L. miodon* (n = 88 scaffolds), scaffolds containing SNPs linked to the putative inversion in *L. miodon* (n = 27 scaffolds), and scaffolds containing SNPs linked to sex in *S. tanganicae* (n = 129 scaffolds). Vertical dashed lines indicate the mean of the distribution for each combination of species and sex, and hash marks along the x-axis show actual data points.

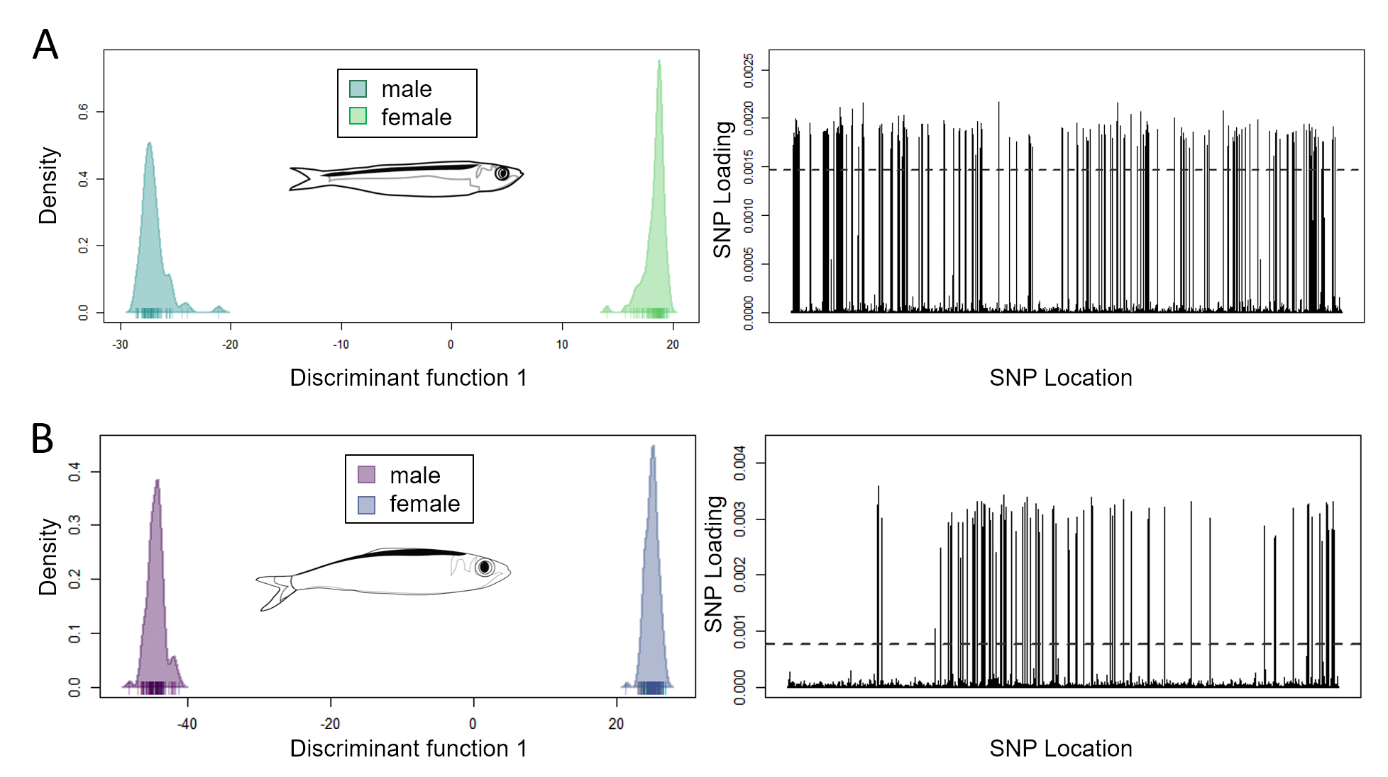

**Figure S6**. DAPC results for differentiation between sexes in species-specific datasets. (A) *S. tanganicae.* Plots show distribution of individuals on discriminant axis 1 (left) and loadings for all SNPs (right, scaffolds ordered from largest to smallest), with threshold indicated delineating 502 significant SNPs (loading > 0.0015) across 129 scaffolds. (B) *L. miodon*, with plots showing distribution of individuals on discriminant axis 1 (left), and loadings for all SNPs (right), with significance threshold indicated delineating 312 significant SNPs (loading > 0.00077) across 88 scaffolds.

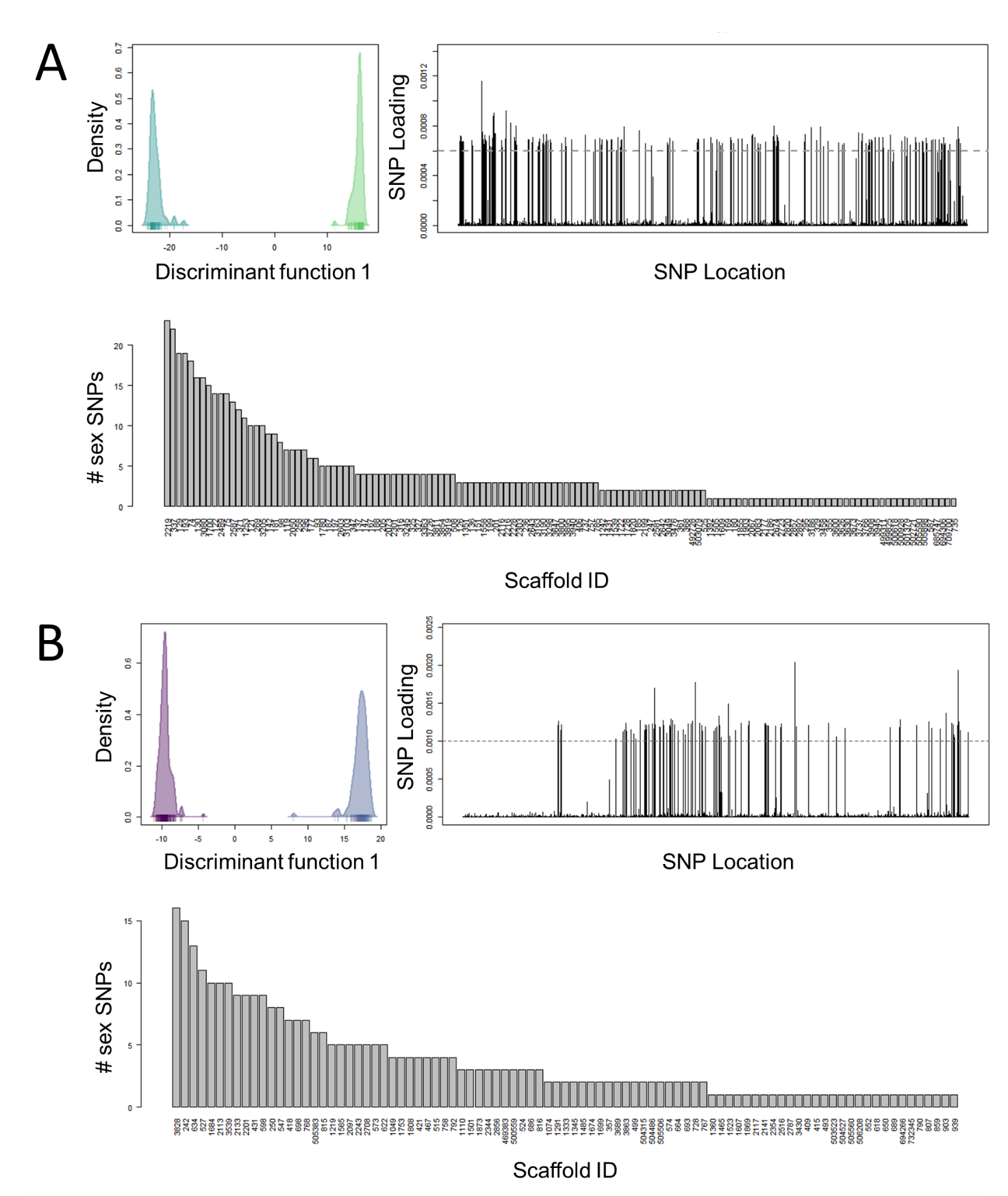

**Figure S7**. DAPC results for sex differentiation in (A) *S. tanganicae* and (B) in *L. miodon* individuals using the combined species SNP dataset. Scaffolds in loading plot are ranked from longest to shortest, but are ranked from most to least sex SNPs in the barplot.

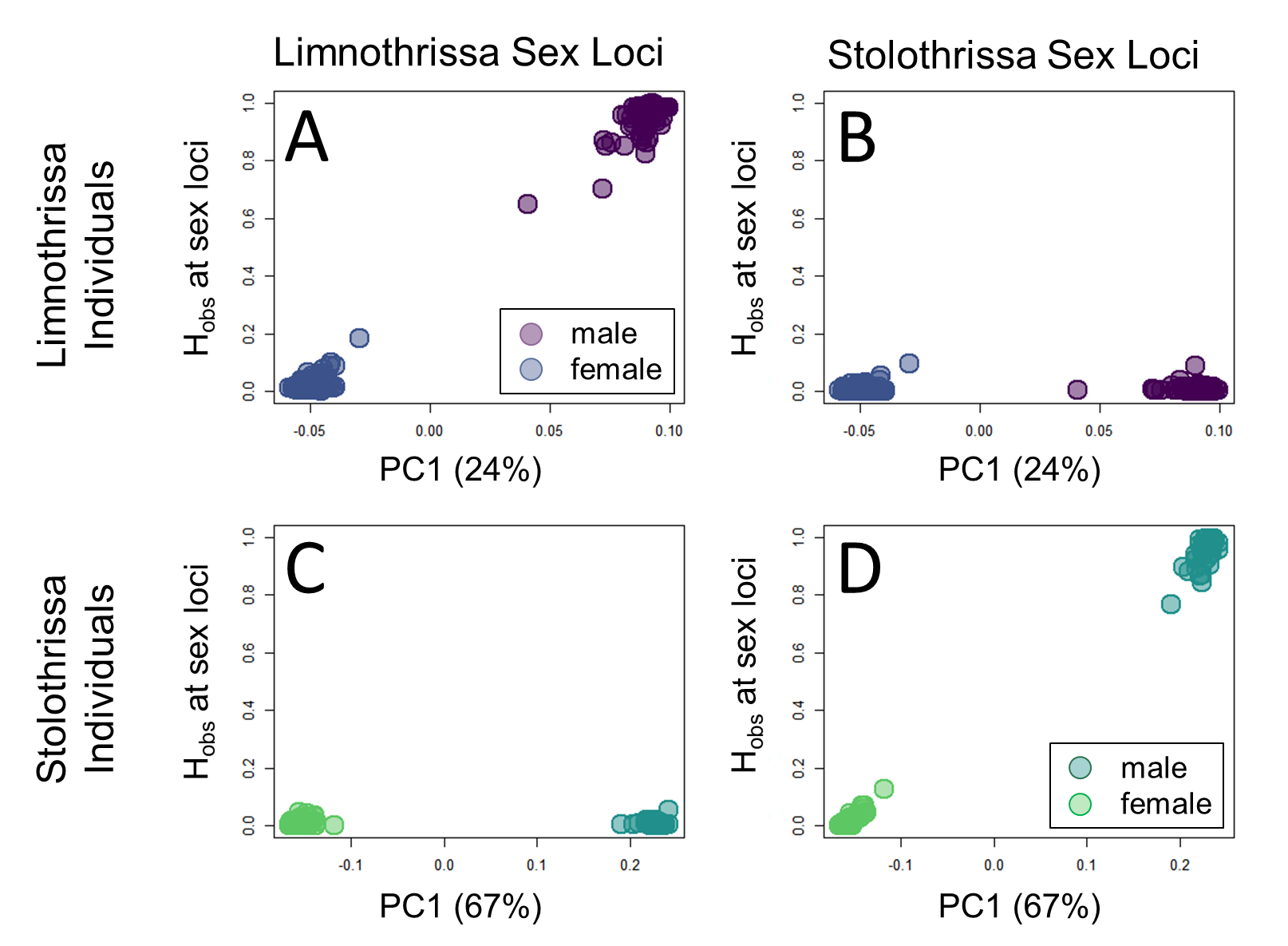

**Figure S8**. Observed heterozygosity of *L. miodon* individuals (A,B) and *S. tanganicae* individuals (C,D) at significant sex loci identified in *L. miodon* (A,C) and *S. tanganicae* (B,D), plotted against the first PC-axis for the species that the individuals belong to. Points are colored by genetically-identified sex, and PCAs were conducted on each species separately using the set of SNPs identified in the species-combined data set.

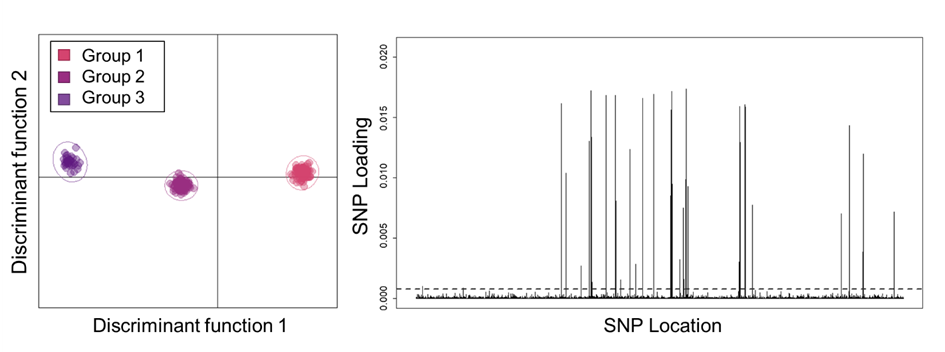

**Figure S9.** DAPC results for *L. miodon* differentiation between groups, showing distribution of locations on discriminant axis 1 and 2 (left), and loadings for all SNPs on discriminant function 1 (right; scaffolds ordered from largest to smallest). We used a cutoff of 0.00077 to identify SNPs with a significant loading on the differences between groups 1 and 3, resulting in a total of 91 significant SNPs across 27 scaffolds.

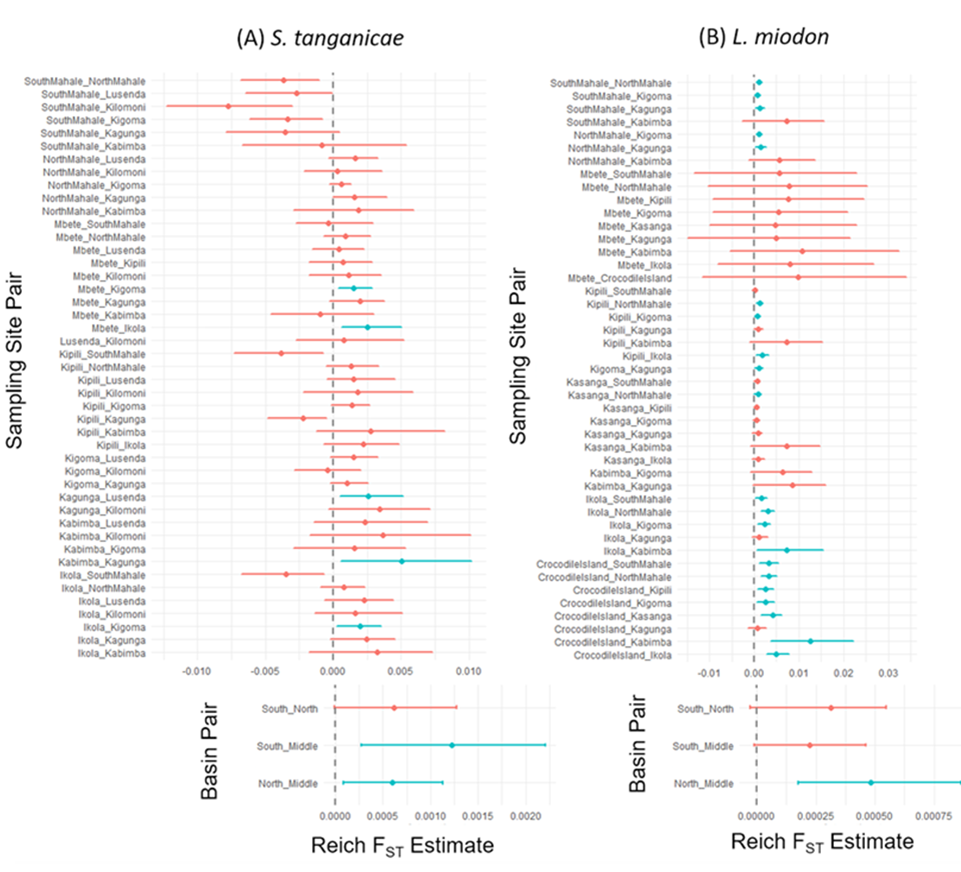

**Figure S10**. Reich F_ST_ point estimates and bootstrapping 95% confidence intervals for each site pair and basin pair of Lake Tanganyika for (A) *S. tanganicae* and (B) *L. miodon*. Color shows significance, with red indicating estimates whose bootstrap confidence interval overlaps 0.

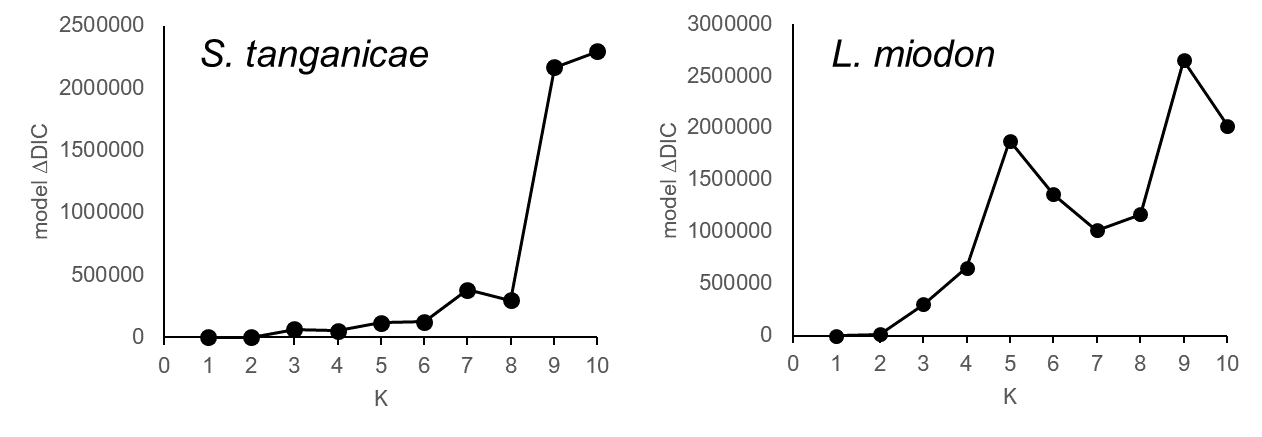

**Figure S11**. Change in deviance information criterion (DIC) for runs of entropy (Gompert et al. 2014) from K=1 to K=10 for *S. tanganicae* and *L. miodon.* Lower DIC values indicate better model fit, and the best fitting model for both species was K=1.

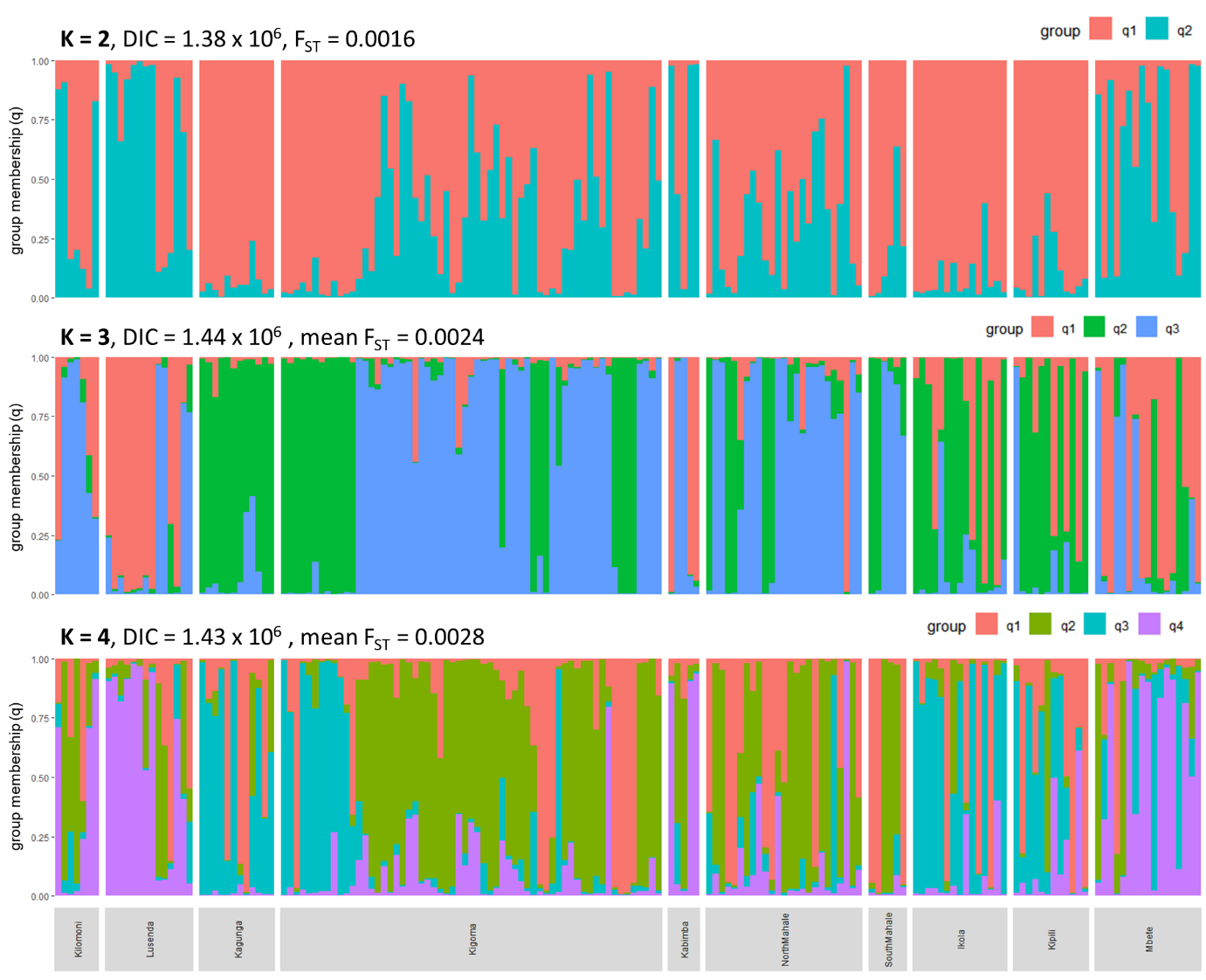

**Figure S12**. Individual group membership (q) based on entropy (Gompert et al. 2014) runs for *S. tanganicae* individuals at K=2, K=3, and K=4 after all sex-linked loci were removed. Samples are separated by sampling locality, indicated across the bottom of each plot. The proportion of ancestry from each cluster is shown by the height of each block of color, with each vertical bar representing a single individual.

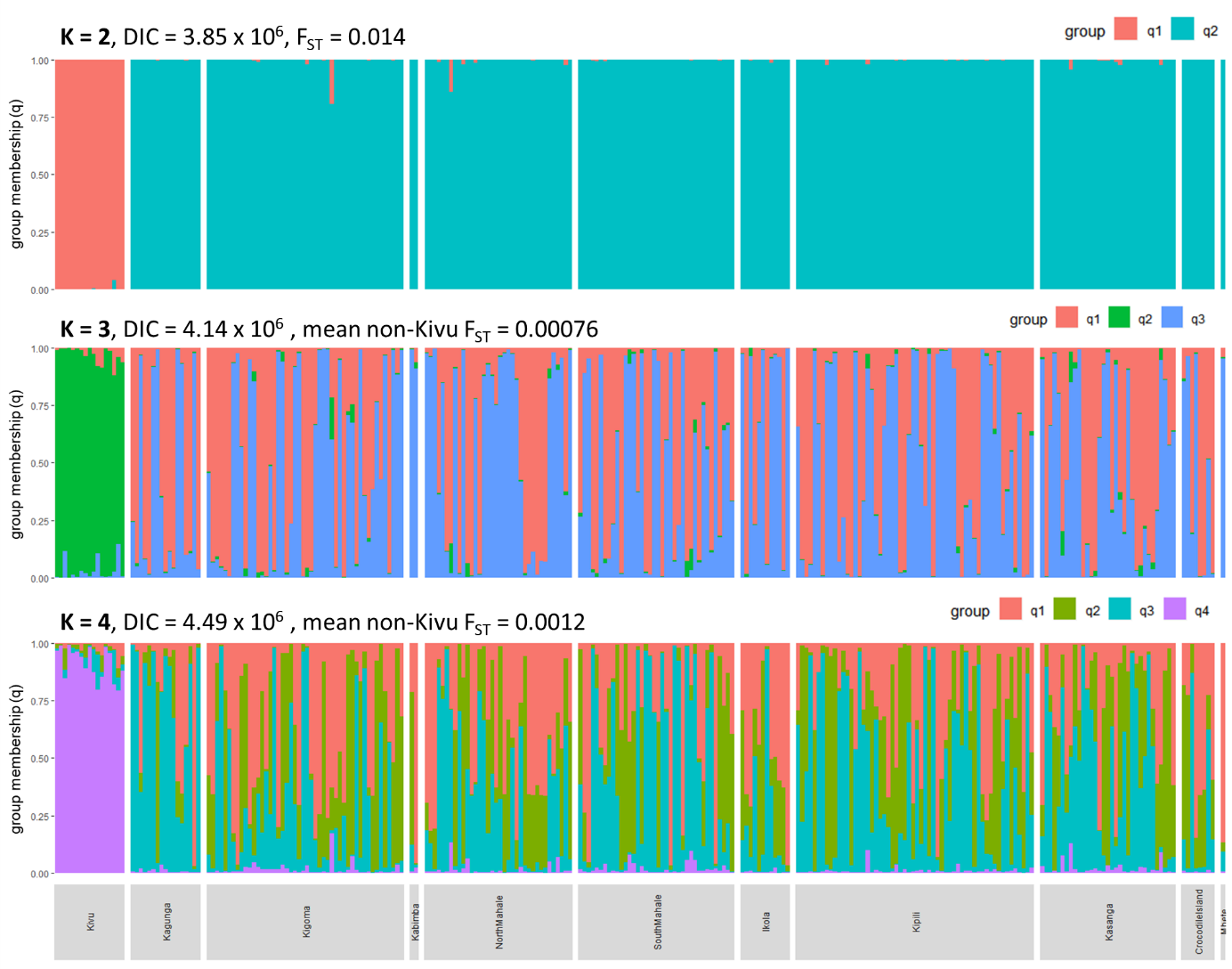

**Figure S13**. Individual group membership (q) based on entropy (Gompert et al. 2014) runs for *L. miodon* individuals at K=2, K=3, and K=4 after all sex-linked loci and the loci linked to a putative inversion were removed. Samples are separated by sampling locations, indicated across the bottom of each plot. The proportion of ancestry from each cluster is shown by the height of each block of color, with each vertical bar representing a single individual.

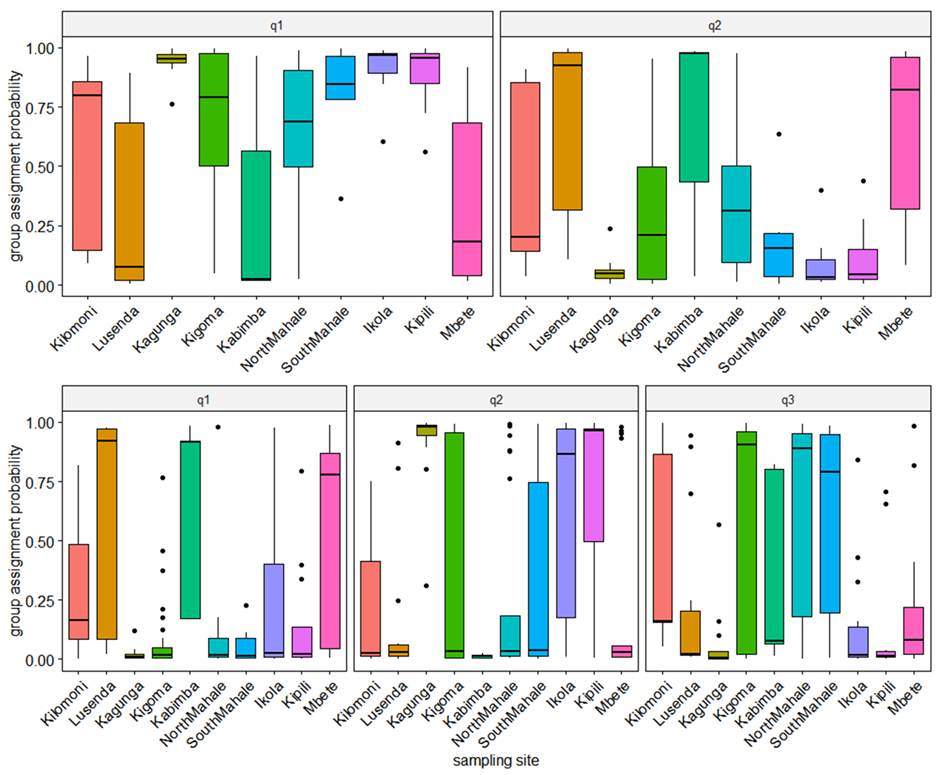

**Figure S14.** Boxplot of group assignment probabilities from entropy models for *S. tanganicae* individuals at K = 2 (top) and K = 3 (bottom), separated by sampling site. In models with K > 1, group assignment probabilities for all groups differed significantly among sampling sites (ANOVA, *p* < 0.001).

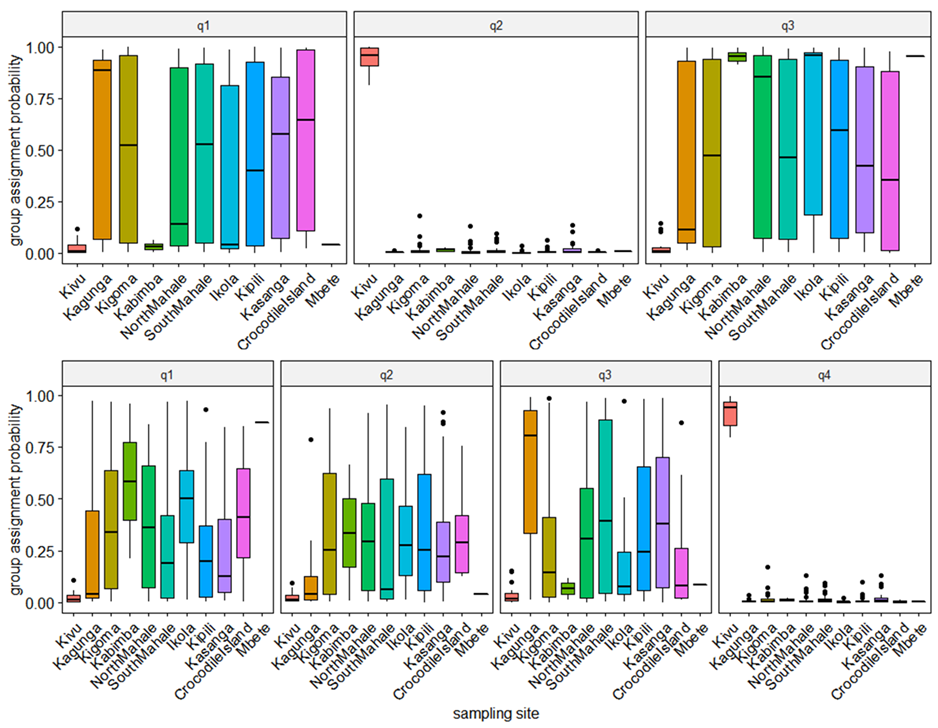

**Figure S15**. Boxplot of group assignment probabilities from entropy models for *L. miodon* individuals at K = 3 (top) and K = 4 (bottom), separated by sampling site. In all models with K > 1, individuals collected in Lake Kivu were a distinct group from the Lake Tanganyika individuals. The non-Kivu groups differed significantly in frequency among Tanganyika sampling sites with more than 2 individuals (ANOVA, p < 0.01)

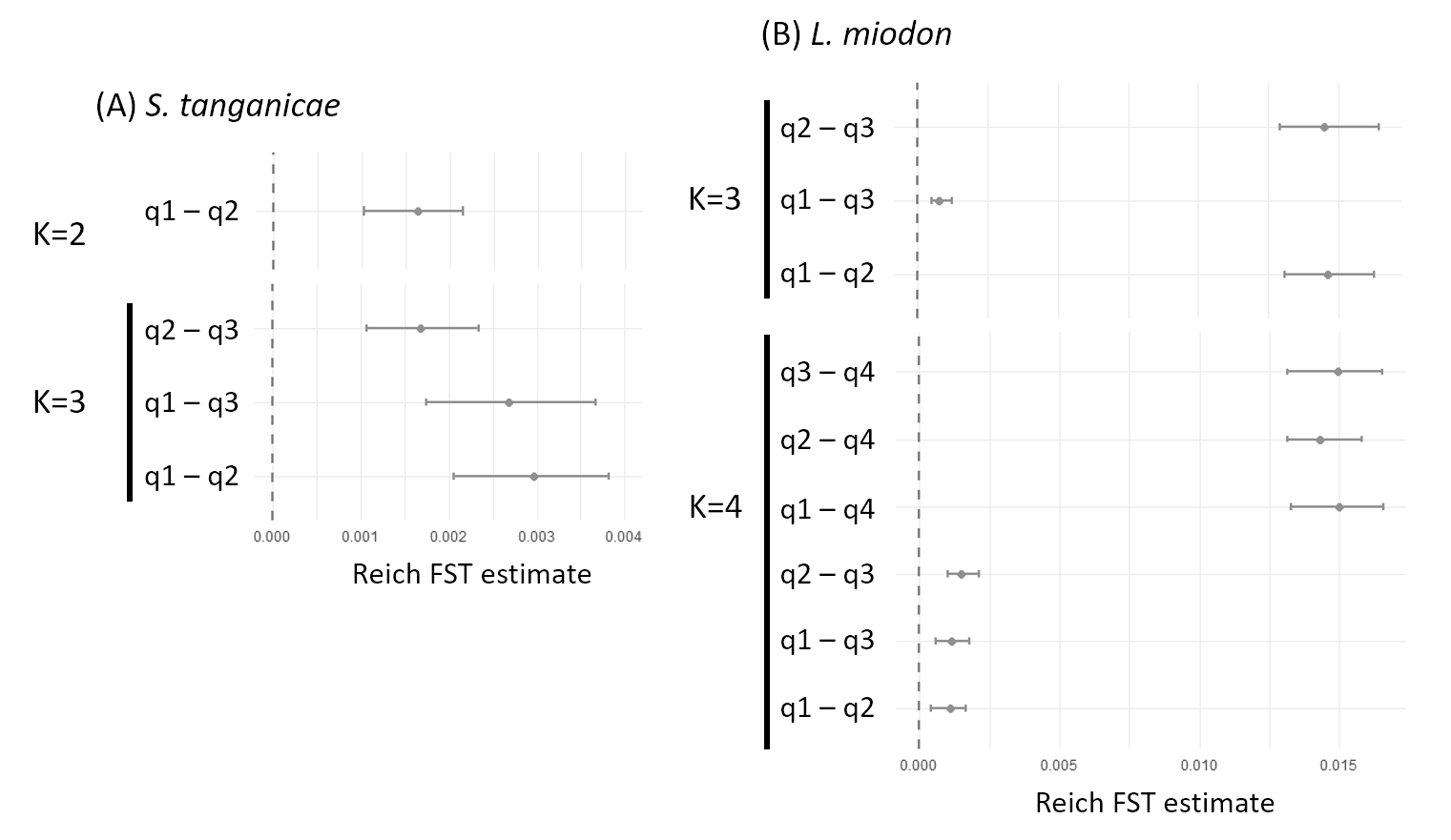

**Figure S16**. Reich F_ST_ point estimates and bootstrapping 95% confidence intervals for each pair of groups identified in entropy for (A) *S. tanganicae* (at K=2 and K=3) and (B) *L. miodon* (at K=3 and K=4). All bootstrap confidence intervals do not overlap with 0.

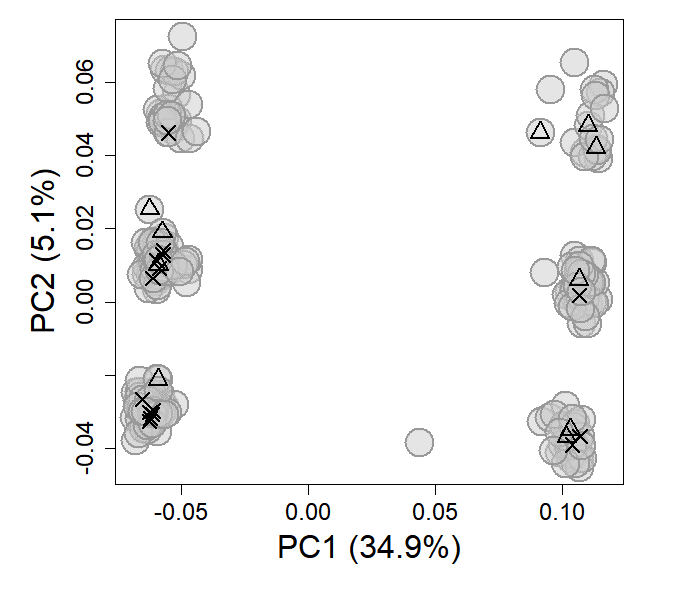

**Figure S17.** PCA of *L. miodon* individuals, highlighting a group of juvenile *L. miodon* (< 3cm) caught from the same school in Sibwesa (South Mahale) in 2015 (black X’s), and nine *L. miodon* fry (< 2cm) caught in one scoop with a hand net in Kagunga in 2017 (black outline triangles). Both single-school samples included individuals from multiple different karyotypes.
